# Dynamics of Cas10 Govern Discrimination between Self and Nonself in Type III CRISPR-Cas Immunity

**DOI:** 10.1101/369744

**Authors:** Ling Wang, Charlie Y. Mo, Michael R. Wasserman, Jakob T. Rostøl, Luciano A. Marraffini, Shixin Liu

## Abstract

Adaptive immune systems are required to accurately distinguish between self and nonself in order to defend against invading pathogens while avoiding autoimmunity. Type III CRISPR-Cas systems employ guide RNAs that recognize complementary RNA molecules to trigger the degradation of both the target transcript and its template DNA. These systems can broadly eliminate foreign targets with multiple mutations, but still effectively curb immunity against the host. The molecular basis for these unique features remains unknown. Here we use single-molecule fluorescence microscopy to study the interaction between a type III-A ribonucleoprotein complex and various RNA substrates. We find that Cas10—the DNase effector of the complex—displays rapid conformational fluctuations on foreign RNA targets, but is locked in a static configuration on self RNA. Single-stranded DNA promotes Cas10’s occupancy at a selected set of conformational states, which is also sensitively modulated by target mutations and predictive of CRISPR interference activity. These findings highlight the central role of the internal dynamics of CRISPR-Cas complexes in self/nonself discrimination and target specificity.

## INTRODUCTION

A fundamental attribute of immune systems is their ability to distinguish foreign from self elements, which is imperative for the host to eliminate invading pathogens while avoiding autoimmunity (Boehm, 2006). Clustered Regularly Interspaced Short Palindromic Repeats (CRISPR) loci and CRISPR-associated (*cas*) genes represent an adaptive immune mechanism for prokaryotes to defend against phage and plasmid infection (Barrangou and Marraffini, 2014; van der Oost et al., 2014). In this mechanism, fragments of the invading DNA are inserted between CRISPR repeats in the host genome. The inserts, known as spacers, are subsequently transcribed and processed into CRISPR RNAs (crRNAs), which assemble with a specific set of Cas proteins to form ribonucleoprotein effector complexes. Upon re-infection, immunity is conferred by crRNA-guided recognition and degradation of the invading genetic element by the effector complex.

Based on their *cas* gene content, CRISPR-Cas systems can be classified into six major types (I-VI) (Koonin et al., 2017). Type III systems, which are identified by their signature *cas10* gene, are further divided into subtypes: III-A/D that encodes the Cas10-Csm complex and III-B/C that encodes the Cas10-Cmr complex. Type III effector complexes employ a uniquely elaborate targeting mechanism (Pyenson and Marraffini, 2017; Tamulaitis et al., 2017), in which active transcription of the target sequence is required for CRISPR immunity (Deng et al., 2013; Goldberg et al., 2014). crRNA derived from the spacer-repeat array guides the Cas10-Csm/Cmr complex to transcribed target RNA containing a protospacer sequence complementary to the crRNA spacer. Multiple copies of the Csm3/Cmr4 subunit in the complex—harboring crRNA-guided RNase activity—cleave the target RNA in 6-nucleotide (nt) intervals (Hale et al., 2009; Samai et al., 2015; Staals et al., 2013; Tamulaitis et al., 2014; Zhang et al., 2012). Binding of the complex to the target RNA further triggers single-stranded DNA (ssDNA) degradation, which is carried out by the Cas10 subunit (Elmore et al., 2016; Estrella et al., 2016; Kazlauskiene et al., 2016). In addition, type III-A loci also encode for Csm6, a nonspecific RNase that becomes essential for immunity when the target is located in late-expressed genes or contains mismatches to the spacer (Jiang et al., 2016).

Such an RNA/DNA dual targeting mechanism contrasts the one employed by type I and II systems, which generally target double-stranded DNA (dsDNA). Moreover, type III systems adopt a distinctive mechanism for self/nonself discrimination. To specify a target, type I and II systems recognize within the invading DNA short (2-4 nt) protospacer adjacent motifs (PAMs), which are absent from the host’s own CRISPR repeats (Gasiunas et al., 2012; Mojica et al., 2009). By contrast, type III systems rely on the crRNA “tag”, an 8-nt sequence derived from the CRISPR repeat located at the 5’ flank of the mature crRNA. Non-complementarity between the crRNA tag and the 3’ flanking sequence of the protospacer licenses a foreign target and triggers an immune response, whereas complementarity specifies host nucleic acids and prevents self targeting (Marraffini and Sontheimer, 2010). Notably, homology between the crRNA tag and the 3’-flanking target sequence does not affect RNA cleavage, but rather inhibits ssDNA cleavage by Cas10 (Kazlauskiene et al., 2016; Samai et al., 2015). However, the molecular mechanism by which Cas10’s DNase activity is switched on or off by the 3’-flanking sequence remains unknown.

Compared to other CRISPR types, type III systems also display an unusually high level of tolerance to mutations in the protospacer sequence (Goldberg et al., 2014; Kazlauskiene et al., 2016; Manica et al., 2013; Maniv et al., 2016; Peng et al., 2015; Staals et al., 2014). A recent comprehensive mutational survey confirmed the broad target specificity and further showed that the accumulation of mutations may weaken, but not abrogate, the immune response to varying degrees depending on the position of the mutations in the protospacer (Pyenson et al., 2017). Nonetheless, how the strength of immunity is differentially modulated by target mutations is still poorly understood.

Single-molecule techniques are powerful tools for dissecting complex protein-nucleic acid interactions and have been applied to study type I and II CRISPR-Cas systems (Blosser et al., 2015; Dagdas et al., 2017; Josephs et al., 2015; Jung et al., 2017; Lim et al., 2016; Loeff et al., 2018; Redding et al., 2015; Singh et al., 2016; Sternberg et al., 2014; Szczelkun et al., 2014; Yang et al., 2018). Here, we used single-molecule fluorescence microscopy to investigate the targeting mechanism of a type III-A Cas10-Csm complex. We found that Cas10 displays strikingly distinct behaviors on self versus nonself RNA: it is locked in a static configuration on host CRISPR transcripts, but samples a large conformational space upon binding to foreign RNA. Among the many states explored by Cas10 on target RNA, a subset is enriched by the presence of ssDNA and sensitively modulated by mutations in the protospacer region of the target. The occupancy of Cas10 at these states is predictive of the CRISPR interference efficiency measured *in vivo*, suggesting that they correspond to the active configuration of the effector complex. These results highlight the exquisite allosteric regulation of the conformational fluctuations of the effector complex by the target sequence, and provide the molecular basis for self/nonself discrimination and mutation tolerance in type III CRISPR-Cas immunity.

## RESULTS

### Single-Molecule Fluorescence Platform for Studying Type III CRISPR-Cas Immunity

We chose the type III-A CRISPR-Cas system from *Staphylococcus epidermidis* as our model system. Its CRISPR loci encode for a Cas10-Csm complex composed of Cas10(×1), Csm2(×3), Csm3(×5), Csm4(×1), Csm5(×1) and a crRNA. We used an engineered *S. epidermidis* CRISPR-Cas locus that contains one single spacer targeting the capsid gene *gp43* of the staphylococcal lytic phage ΦNM1γ6 (Figures S1A) (Jiang et al., 2016). Cas10-Csm complexes harboring mature crRNA were heterologously expressed in *Escherichia coli* (Figure S1B). For single-molecule experiments, RNA substrates were labeled with a biotin and a Cy3 fluorophore at opposite ends. Individual RNA molecules were immobilized on a glass coverslip and their fluorescence signals were detected by total-internal-reflection fluorescence microscopy. RNA cleavage by the Cas10-Csm complex would result in release of the fluorophore into the solution and, thus, a decrease in the surface density of Cy3 fluorescent spots (Figure 1A). Three types of RNA substrates were assessed (Figure 1B): (1) a wildtype (“WT”) RNA— mimicking *bona fide* RNA targets—that contains a 35-nt protospacer sequence complementary to the crRNA spacer and a 3’-flanking sequence that is non-complementary to the crRNA 5’ tag; (2) an “Anti-tag” RNA—mimicking RNA molecules generated by antisense transcription of the host’s own CRISPR array—that contains both a matching protospacer sequence and an 8-nt 3’-flanking anti-tag sequence; and (3) a “Non-specific” RNA containing a scrambled sequence with no homology to the crRNA. In the presence of Mg^2+^, the surface densities of WT and Anti-tag RNAs decreased at similar rates, demonstrating that they were both efficiently cleaved by Csm3 (Figures 1C and 1D). The rate obtained from the single-molecule assay was comparable to that measured in bulk (Figure 1D). In contrast, minimal cleavage was observed with the Non-specific RNA or in the absence of Mg^2+^ (Figures 1D and S2). These results are consistent with previous studies showing that base pairing with the crRNA tag, as is the case for the Anti-tag RNA, does not inhibit the RNA cleavage activity of the effector complex (Samai et al., 2015; Tamulaitis et al., 2014). It is noteworthy that RNA cleavage products are quickly released by the complex—as reflected by the disappearance of fluorescence from the surface—unlike Cas9-mediated DNA targeting in which DNA remains stably bound to the complex even after cleavage (Sternberg et al., 2014). This feature potentially allows the type III effector complex to process multiple targets within a short time window.

**Figure 1.**
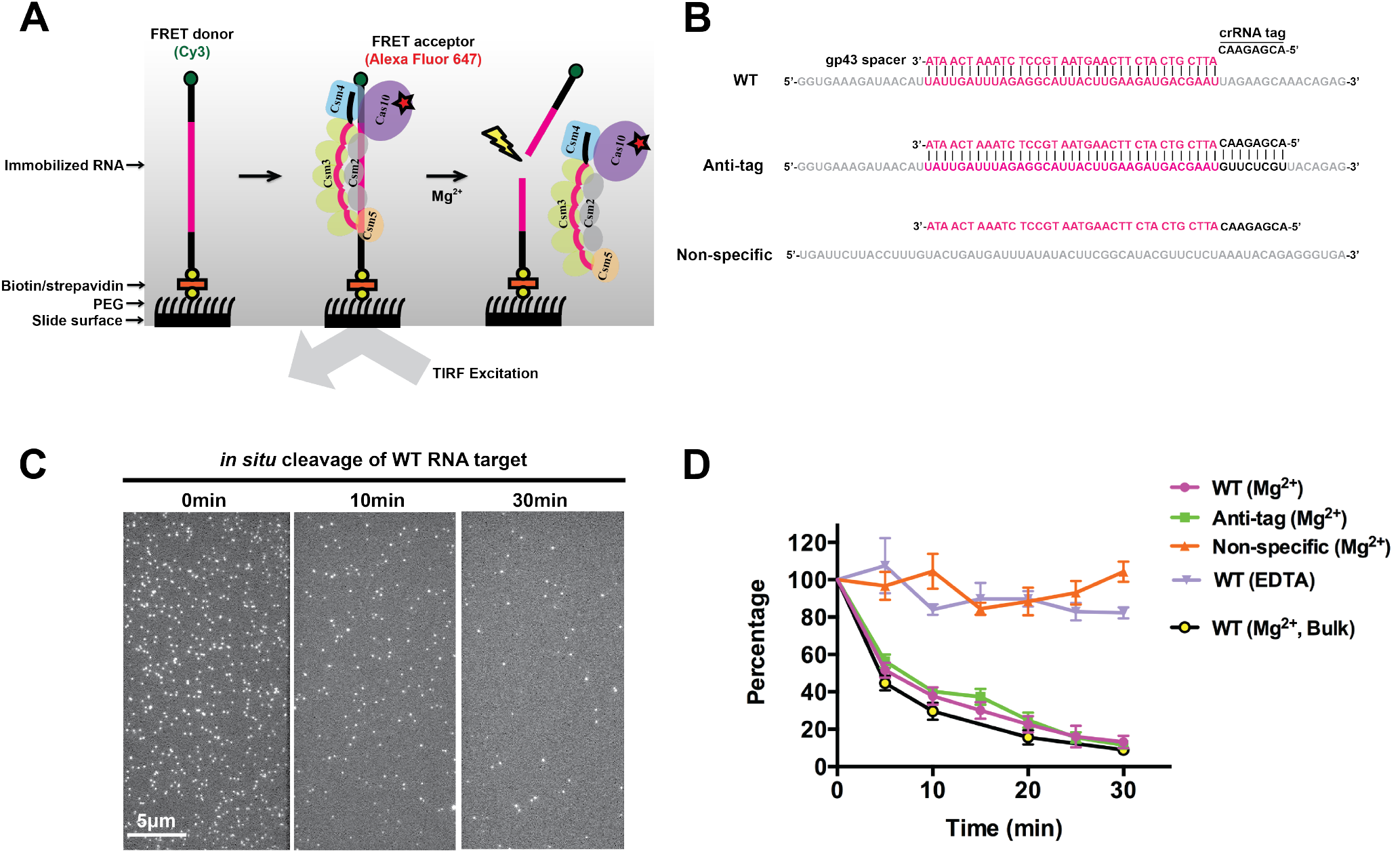
A Single-Molecule Fluorescence Platform for Studying Type III CRISPR-Cas Targeting Mechanism. (**A**) Schematic of the single-molecule fluorescence imaging platform. (**B**) Sequences of the WT, Anti-tag and Non-specific RNAs. (**C**) Representative fields of view on a total-internal-reflection fluorescence microscope showing surface-immobilized Cy3-labeled RNA molecules. Disappearance of the fluorescent spots after Mg^2+^ addition reflects the cleavage of individual RNA molecules. (**D**) Cleavage kinetics measured for different RNA substrates from the single-molecule assay, which are plotted as the average surface density of molecules versus time after Mg^2+^ addition. The surface density of WT RNA remained unchanged when an EDTA-containing buffer without Mg^2+^ was added (purple triangles), demonstrating minimal photobleaching during the time of observation. The WT RNA cleavage kinetics measured from a bulk assay is shown in yellow circles. Data are represented as mean ± SD from multiple fields of view (*N* > 10) for the single-molecule assay or three independent experiments for the bulk assay.

### Dynamic Interaction Between the Cas10-Csm Complex and Its RNA Target

Since both self and nonself RNAs can be equally recognized by type III CRISPR complexes for cleavage, we postulated that the discrimination may be conferred by distinct binding configurations of the Cas10-Csm complex on different RNA targets. To test this hypothesis, we used single-molecule fluorescence resonance energy transfer (FRET) to probe the interactions between Cas10-Csm and various RNA substrates. We began by focusing on the WT RNA, which mimics transcripts of foreign elements. We designed several FRET labeling schemes based on the structural model for the target-bound Cas10-Csm complex (Kazlauskiene et al., 2016; Osawa et al., 2015; Tamulaitis et al., 2017; Taylor et al., 2015). First, we attached the FRET donor (Cy3) to the 5’ end of the WT RNA and the FRET acceptor (AlexaFluor647) to the Csm5 subunit, which is located at the distal side of the crRNA tag and Cas10 (Figure 2A). Single-molecule data were collected in an EDTA-containing buffer in order to prevent RNA degradation. Binding of Csm5-labeled Cas10-Csm complexes to 5’-end-labeled WT RNA resulted in a stable FRET state (Figure 2A). The distribution of FRET efficiency (*E*) built from many molecules displayed a single peak centered at ~ 0.25 (Figure 2B). This result suggests that the Cas10-distal end of the target-bound complex is largely static.

**Figure 2.**
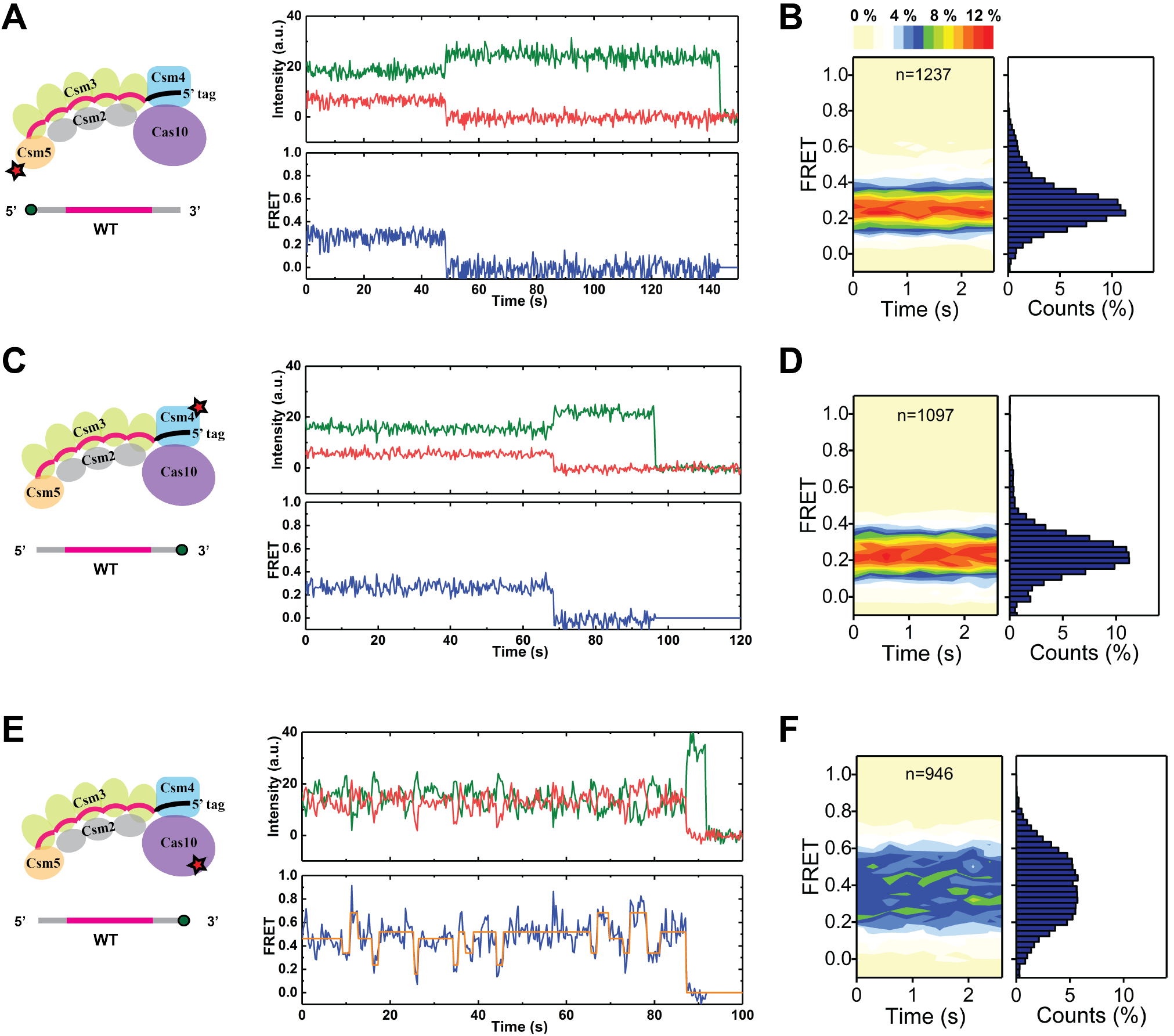
Interaction between the Cas10-Csm Complex and Nonself RNA. (**A**) A representative time trajectory of donor (Cy3, green) and acceptor (AlexaFluor647, red) fluorescence intensities and the corresponding FRET values (blue) using acceptor-labeled Csm5 subunit and donor-labeled WT RNA at its 5’-end. (**B**) Contour plot and histogram for the FRET distribution from single-molecule trajectories described in (A) (*N* = 1237; *n* denotes the number of molecules analyzed). (**C** and **D**) A representative fluorescence and FRET trajectory (C) and the corresponding FRET contour plot and histogram (D) using acceptor-labeled Csm4 subunit and donor-labeled WT RNA at its 3’-end (*N* = 1097). (**E** and **F**) A representative fluorescence and FRET trajectory (E) and the corresponding FRET contour plot and histogram (F) using acceptor-labeled Cas10 subunit and donor-labeled WT RNA at its 3’-end (*N* = 946). The orange line plots idealized FRET states from hidden-Markov-modeling analysis.

We then moved the FRET pair to the Cas10-proximal end of the complex, with the donor attached to the 3’ end of the WT RNA and the acceptor labeled on the Csm4 subunit that makes contacts with the crRNA tag (Figure 2C). Again we observed one predominant FRET state (*E* ~ 0.23) (Figures 2D), suggesting that the 3’-flanking region of the WT RNA, even though unable to base pair with the crRNA tag, is nonetheless stationary relative to Csm4.

Finally, we placed the acceptor fluorophore on Cas10—the largest subunit in the complex that harbors the DNase activity—while keeping the donor at the 3’ end of the WT RNA (Figure 2E). In contrast to the previous two labeling schemes, most of the FRET trajectories obtained with Cas10-labeled complexes were highly dynamic, rapidly sampling many different states (Figures 2E and S3). Accordingly, we observed a broad FRET distribution with *E* spanning from 0.1 to 0.8 (Figure 2F). This finding reveals that Cas10, the signature protein of all type III systems, is highly mobile relative to the rest of the complex when bound to nonself RNA targets.

### Distinct Behaviors of Cas10 on Self Versus Nonself RNA

Next we performed the same set of FRET measurements on the Anti-tag RNA, which mimics transcripts derived from the host’s own CRISPR loci. The FRET distribution for Csm5-labeled complexes on 5’-labeled Anti-tag RNA showed a single peak at ~ 0.25 (Figures 3A and 3B), indistinguishable from that for the WT RNA (Figure 2B). The FRET distribution for Csm4-labeled complexes on 3’-labeled Anti-tag RNA again showed a single peak (Figures 3C and 3D), but with a modest increase in the FRET value (*E* ~ 0.28) compared to the corresponding distribution for the WT RNA (Figure 2D). This difference can be rationalized by the base pairing between the crRNA tag and the 3’-flanking region of the Anti-tag RNA, which conceivably brings the 3’ end of the RNA closer to Csm4 (Tamulaitis et al., 2017).

**Figure 3.**
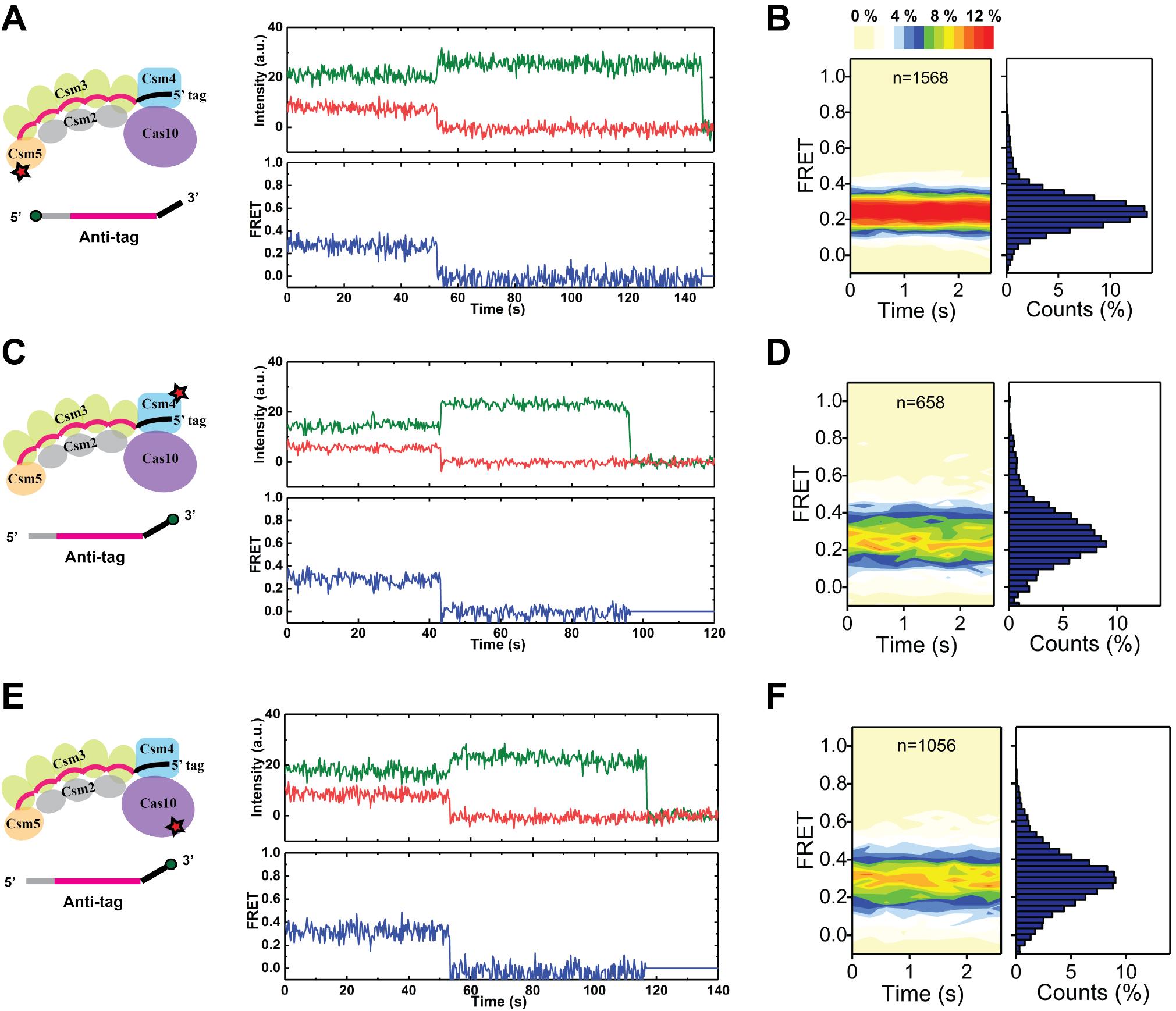
Interaction between the Cas10-Csm Complex and Self RNA. (**A**) A representative time trajectory of fluorescence intensities and FRET values using acceptor-labeled Csm5 subunit and donor-labeled Anti-tag RNA at its 5’-end. (**B**) Contour plot and histogram for the FRET distribution from single-molecule trajectories described in (A) (*N* = 1568). (**C** and **D**) A representative fluorescence and FRET trajectory (C) and the corresponding FRET contour plot and histogram (D) using acceptor-labeled Csm4 subunit and donor-labeled Anti-tag RNA at its 3’-end (*N* = 658). (**E** and **F**) A representative fluorescence and FRET trajectory (E) and the corresponding FRET contour plot and histogram (F) using acceptor-labeled Cas10 subunit and donor-labeled Anti-tag RNA at its 3’-end (*N* = 1056).

Strikingly, interrogation of Cas10-labeled complexes on 3’-labeled Anti-tag RNA revealed a major difference. The vast majority of Cas10 binding events on the Anti-tag RNA exhibited a stable, low FRET state (*E* ~ 0.29) (Figures 3E and 3F), in stark contrast to the wide fluctuations observed in the FRET traces of Cas10 on the WT RNA (Figures 2E and 2F).

### Self RNA Inhibits Activation of the Cas10-Csm Complex

To correlate the single-molecule data to the *in vivo* immune responses elicited by the WT and Anti-tag RNAs, we conducted a bacterial transformation assay developed previously to measure the effect of target mutations on CRISPR immunity (Marraffini and Sontheimer, 2010; Samai et al., 2015), but with the same target sequence used in the single-molecule experiments of the current study. In this assay, *Staphylococcus aureus* RN4220 strains were transformed with two plasmids: (1) pCRISPR carrying an *S. epidermidis* type III-A CRISPR-Cas system with the *gp43* spacer or a control plasmid (pCRISPRΔspc) with a non-matching spacer; and (2) a plasmid encoding either the WT RNA target (pTarget^WT^) or the Anti-tag RNA (pTarget^Anti-tag^) under the control of an anhydrotetracycline (aTc)-inducible promoter (Figure 4A). Activation of Cas10 by target RNA binding would lead to degradation of the target plasmid and inhibition of transformation. In the absence of aTc, there was no target transcription to activate Cas10 and, therefore, no degradation of pTarget DNA. As expected, we measured a high efficiency of transformation for both pTarget^WT^ and pTarget^Anti-tag^ (Figure 4B). In the presence of aTc, transformation of pTarget^WT^ was essentially abrogated, suggesting effective elimination of the plasmid DNA by Cas10. In contrast, pTarget^Anti-tag^ still displayed substantial transformation efficiency indistinguishable from the pCRISPRΔspc control, indicating that the CRISPR immunity is greatly diminished by the Anti-tag RNA (Figures 4B and 4C).

**Figure 4.**
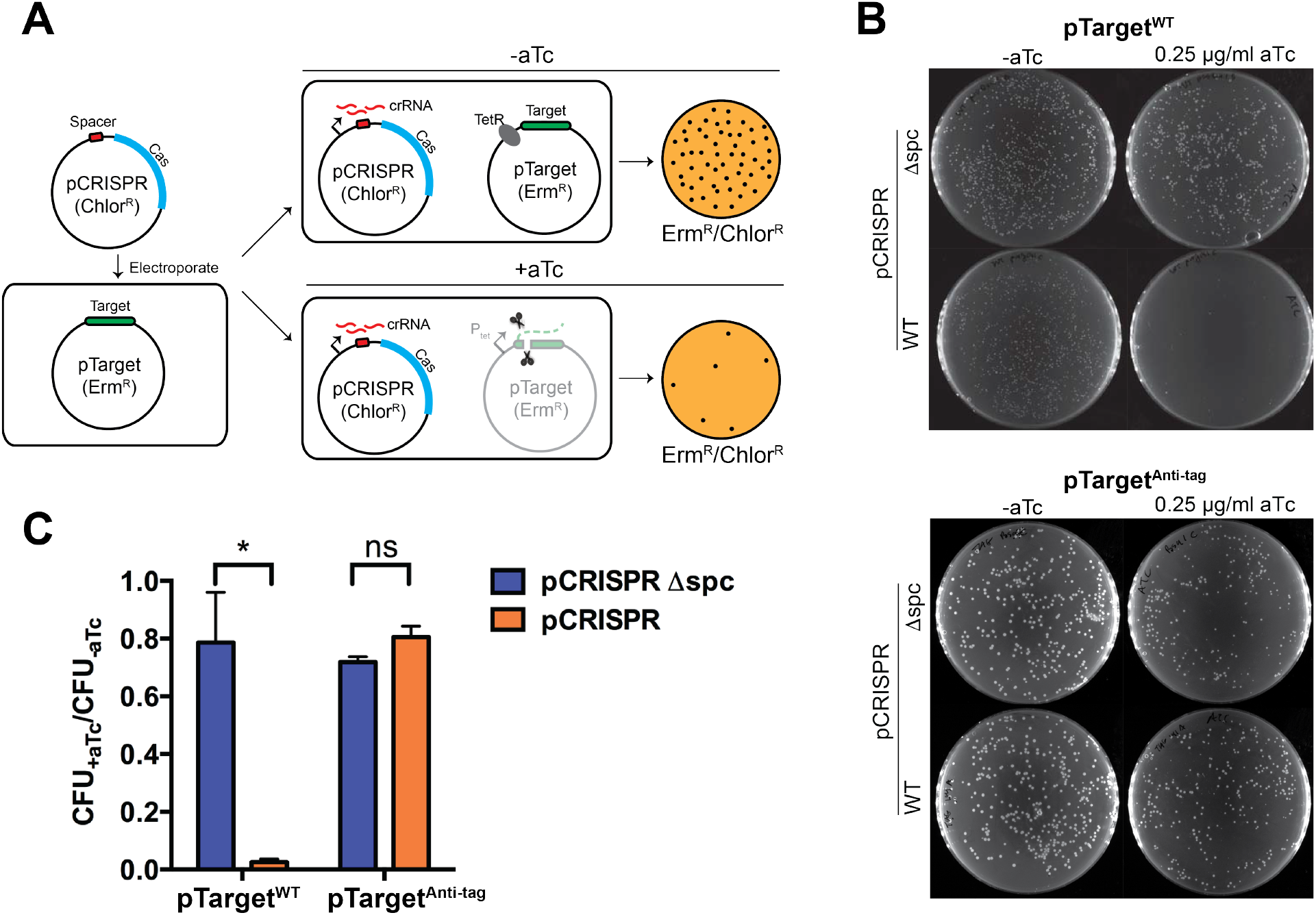
Evaluation of Type III CRISPR Immunity Against Self Versus Nonself Elements. (**A**) Schematic of the bacterial transformation assay. *S. aureus* strains with pTarget (Erm^R^) were transformed with pCRISPR (Chlor^R^) and plated onto chloramphenicol and erythromycin for double selection. In the absence of aTc, the tetracycline repressor (TetR) prevents target transcription and therefore CRISPR immunity against pTarget. In the presence of aTc, the target sequence on pTarget is transcribed, which triggers CRISPR immunity against the plasmid. Degradation of pTarget results in the loss of erythromycin resistance, thereby reducing the transformation efficiency. (**B**) Representative plates of staphylococci colonies under different targeting conditions. In the presence of aTc, transformation of pTarget^WT^ plasmid was greatly diminished, indicating the effective immunity induced by WT RNA transcription. By contrast, pTarget^Anti-tag^ plasmid remained a similar high level of transformation as the non-induction condition, suggesting impaired CRISPR immunity caused by the Anti-tag RNA. As a control, both plasmids had similar high efficiencies of transformation into cells harboring pCRISPRΔspc. (**C**) Transformation efficiencies of pTarget^WT^ and pTarget^Anti-tag^ into cells containing the pCRISPR (orange bars) or pCRISPRΔspc (blue bars) plasmid. The transformation efficiency is calculated as the ratio of colony-forming units (CFU) per microgram of plasmid DNA transformed in the presence and absence of aTc. Data are represented as mean ± SEM (three independent experiments).

The *in vitro* and *in vivo* results together suggest that the distinct behaviors of Cas10 on WT and Anti-tag RNAs are correlated to its ability to provide immunity to the host cell: the stable ~ 0.30 FRET state observed with the Anti-tag RNA likely represents an inactive configuration of Cas10; whereas binding of the WT RNA unlocks Cas10 and prompts it to quickly access many conformational states, a subset of which enables DNase activity to degrade the invader plasmid.

### Protospacer Mutations Differentially Modulate Cas10 Dynamics

We have shown that complementarity between the 3’ flanking region of the protospacer and the crRNA 5’ tag dramatically influences the behavior of Cas10. Next we sought to investigate the effect of mismatches within the protospacer on Cas10 dynamics. We mutagenized the first (closest to the crRNA tag), second, or last 10-nt segments of the 35-nt-long protospacer sequence to create mismatches against the corresponding segment of the crRNA spacer. The mutated RNA targets are termed MM1-10, MM11-20 and MM26-35, respectively (Figure 5A). Bulk biochemical experiments showed that mismatches only inhibited RNA cleavage within the mutated segment but not neighboring segments (Figure S4), confirming the requirement of base pairing between spacer and protospacer for RNA cleavage and the independent activities of the multiple copies of Csm3 (Staals et al., 2014).

**Figure 5.**
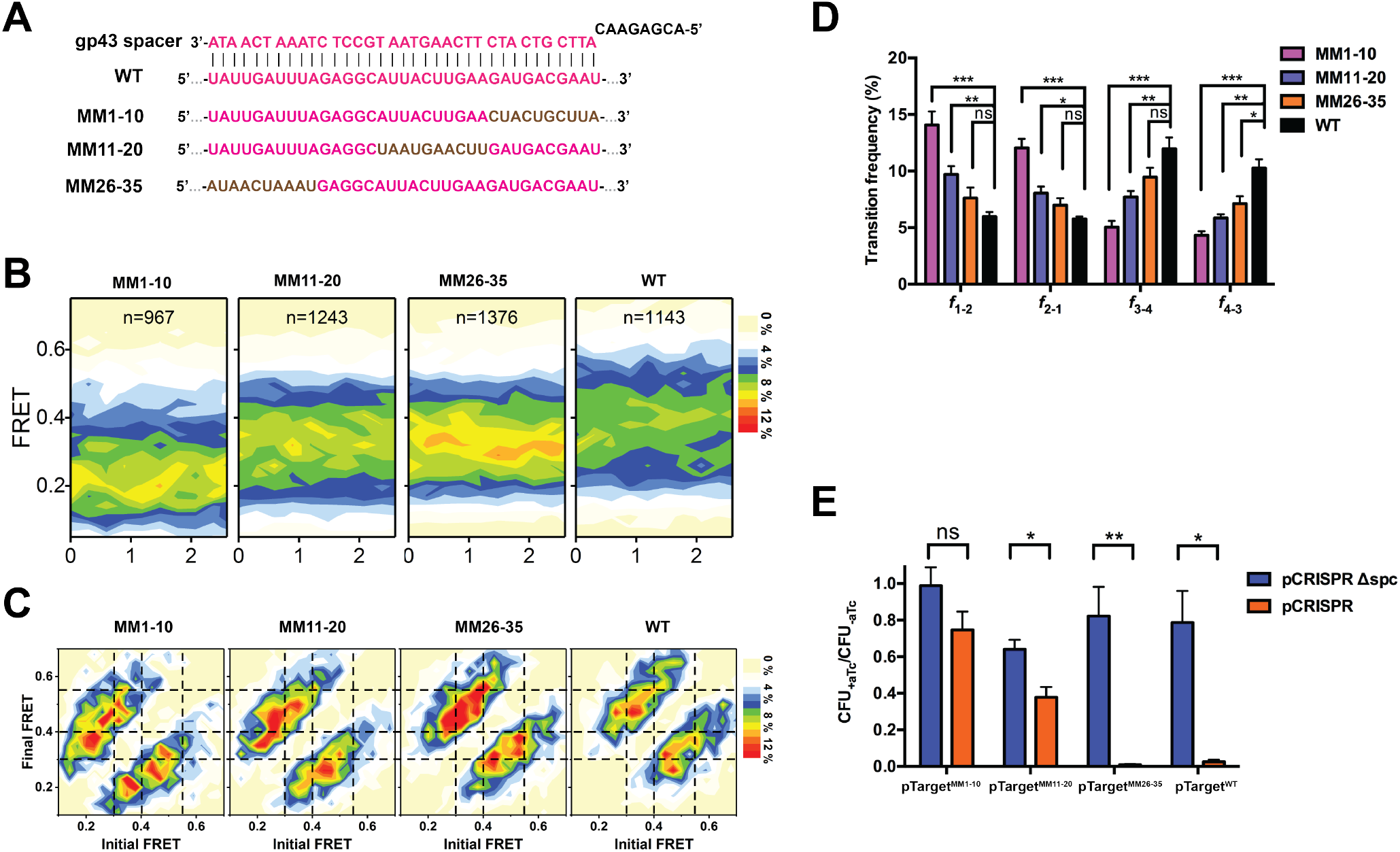
Protospacer Mutations Modulate Cas10 Dynamics and Type III CRISPR Immunity. (**A**) Sequences of different mismatched RNA targets. Mutated regions are shown in brown. (**B**) FRET contour plots for WT and mismatched RNA targets. FRET donor and acceptor were placed on the 3’ end of RNA and Cas10, respectively. (**C**) Transition density plots for WT and mismatched RNA targets. Dashed lines separate distinct FRET groups: G_1_ (*E* ≤ 0.3), G_2_ (0.3 < *E* ≤ 0.4), G_3_ (0.4 < *E* ≤ 0.55) and G_4_ (*E* > 0.55). (**D**) Transition frequencies between FRET groups for different RNA targets. For example, *f*_1-2_ denotes the transition frequency from G_1_ to G_2_. (**E**) Transformation efficiencies of pTarget^WT^, pTarget^MM1-10^, pTarget^MM11-20^ and pTarget^MM26-35^ into cells containing the pCRISPR plasmid (orange bars). A smaller value corresponds to a stronger immune response. The same measurements were repeated using pCRISPRΔspc-containing cells as a negative control (blue bars). Data are represented as mean ± SEM.

We then performed the single-molecule FRET assay to interrogate the interactions of the Cas10-Csm complex with the mismatched RNA targets. We used Cas10-Csm complexes harboring RNase-deficient Csm3^D32A^ mutants in order to monitor the behavior of Cas10 in a Mg^2+^-containing buffer. Cas10 exhibits conformational fluctuations on all mismatched targets (Figure S5A), similar to the WT RNA but in opposition to the Anti-tag RNA. Interestingly, the FRET distribution varied among different targets, shifting toward lower FRET values and deviating further from the WT distribution as the mismatches move from tag-distal to tag-proximal regions (Figure 5B).

To quantify the effects of mismatches on Cas10 dynamics, we employed hidden-Markov-modeling (HMM) analysis (McKinney et al., 2006) to identify distinct FRET states in the single-molecule trajectories and transitions between them (orange lines in Figures 2E, S3, and S5A). The resulting transition density plots (TDP) display the relative frequencies of transitions binned by the FRET values before and after each transition. We separated the HMM-fitted states into four groups: G_1_ (*E* ≤ 0.3), G_2_ (0.3 < *E* ≤ 0.4), G_3_ (0.4 < *E* ≤ 0.55), and G_4_ (*E* > 0.55) (Figure 5C). A comparison of TDPs revealed that the WT, MM26-35, MM11-20 and MM1-10 RNAs exhibit transition frequencies between the high FRET groups G_3_ and G_4_ (*f*_3-4_ and *f*_4-3_) in a descending order (Figures 5D and S5B). The opposite pattern was observed for the transition frequencies between the low FRET groups (*f*_1-2_ and *f*_2-1_) (Figure 5D). These results demonstrate that protospacer mutations differentially influence the conformational distribution of Cas10, with tag-proximal mutations (MM1-10) having the strongest effect and tag-distal ones (MM26-35) the weakest.

### Conformational Distribution of Cas10 Correlates with CRISPR Interference Activity

To investigate the relationship between the conformational distribution of Cas10 and the strength of type III CRISPR immunity, next we performed the transformation assay described in Figure 4A with pTarget plasmids encoding the mismatched RNA targets (pTarget^MM1-10^, pTarget^MM11-20^ and pTarget^MM26-35^). The strength of immunity—reflected by the pTarget transformation efficiency—decreases as the target mismatches move from tag-distal (MM26-35) to tag-proximal (MM1-10) regions (Figure 5E, orange bars). Importantly, this trend matches well with the gradual shift in the distribution of Cas10’s conformational states obtained from the single-molecule FRET measurements (Figures 5B-5D). Hence, the conformational distribution of Cas10 is correlated with *in vivo* CRISPR interference activity: the more time Cas10 spends in the high FRET states, the stronger immunity the Cas10-Csm complex confers.

### Single-Stranded DNA Enriches Specific Cas10 Conformations

Among the various RNA targets studied here, Anti-tag and MM1-10 elicited the weakest immune responses (Figures 4C and 5E). They also resulted in predominantly low FRET populations (*E* < 0.4) upon Cas10-Csm binding (Figures 3F and 5B). On the contrary, WT and MM26-35 RNAs induced Cas10 to occupy higher FRET states and, accordingly, triggered robust anti-plasmid immunity (Figures 5B-5E). These results prompted us to propose that Cas10 becomes DNase-active when accessing the higher FRET states (*E* > 0.4). To further determine the identity of the DNase-active state, we reasoned that it would be enriched by the engagement of DNA substrates. Thus, we examined the effect of DNA on the conformational fluctuations of Cas10.

Using the Cas10/RNA-3’-end labeling scheme (Figure 2E), we obtained single-molecule FRET trajectories with the WT RNA target in the presence of 500 nM linear ssDNA that is 55-nt long and contains the same base sequence as the WT RNA (Figures 6A and S6A). The trajectories remained highly dynamic, but the relative population of the highest FRET group G_4_ (*E* > 0.55) significantly increased (Figure 6B). Moreover, TDP analysis revealed that ssDNA caused a higher probability for effector complexes residing in other FRET groups to transition to G_4_ (Figures 6C and S6B). Interestingly, double-stranded DNA (dsDNA) of the same length and sequence had little effect on the FRET distribution (Figures 6B and 6C), corroborating previous reports that dsDNA is not a good substrate for Cas10 (Estrella et al., 2016; Kazlauskiene et al., 2016). Collectively, these results strongly suggest that G_4_ represents the DNase-active form of the Cas10-Csm effector complex.

**Figure 6.**
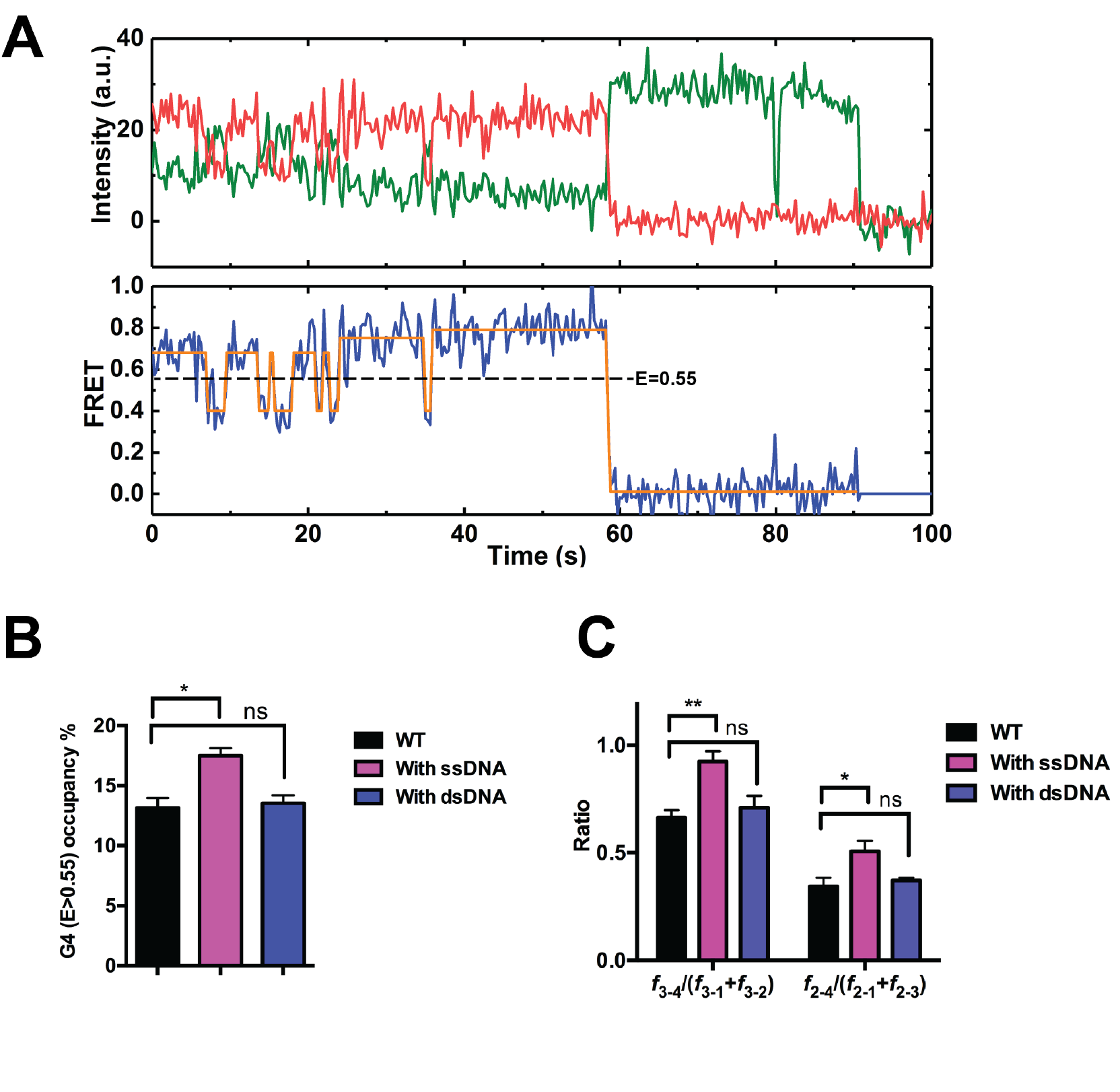
Effects of DNA on Cas10’s Conformational Distribution. (**A**) A representative fluorescence and FRET trajectory using acceptor-labeled Cas10 and donor-labeled WT RNA at its 3’ end in the presence of 500 nM single-stranded DNA. (**B**) The fraction of time the complex spent within the highest FRET group G_4_ (*E* > 0.55) in the absence of DNA (black), in the presence of 500 nM ssDNA (magenta), and in the presence of 500 nM dsDNA (blue). (**C**) The likelihood of complexes in G_2_ or G_3_ transitioning to G_4_—shown as the ratio of transition frequencies to G_4_ and to the other groups—with or without DNA. Data are represented as mean ± SEM.

## DISCUSSION

Type III CRISPR-Cas systems employ an elaborate targeting mechanism to degrade both the invading DNA and its RNA transcripts. The extraordinary complexity compared to other CRISPR systems allows for exquisite spatiotemporal control of the immune response (Tamulaitis et al., 2017). Results presented here for the first time provide a molecular explanation for the discrimination between self and nonself and the unusually high tolerance to target mutations during type III CRISPR immunity (Figure 7). Central to our findings is the remarkable conformational flexibility of Cas10—the signature subunit of type III effector complexes. We show that the conformational distribution of Cas10 plays a determining role in the CRISPR interference activity of the effector complex. Interestingly, the dynamic nature of the complex appears to be modular—the other non-catalytic scaffold subunits, such as Csm4 and Csm5, are largely immobile relative to the RNA target.

**Figure7.**
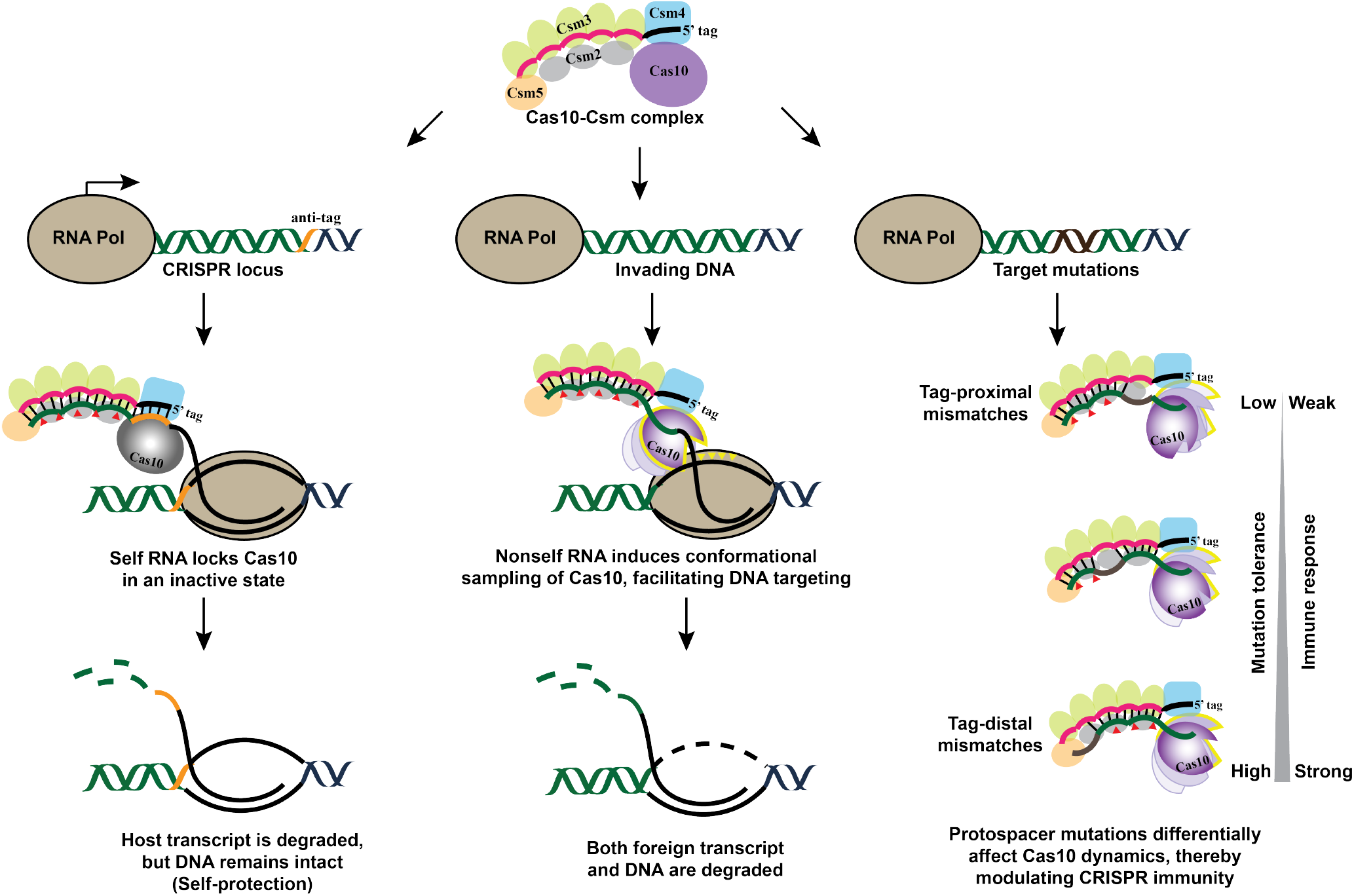
Model for Self/Nonself Discrimination and Target Mutation Tolerance in Type III CRISPR-Cas Immunity. Complementarity between the crRNA 5’ tag and the anti-tag sequence within the RNA transcribed from the host’s CRISPR locus locks Cas10 in an inactive state and suppresses CRISPR immunity. On the contrary, trology to the crRNA tag, thereby enabling conformational fluctuations of Cas10 and conferring robust immunity. Mutations in the protospacer sequence of the target differentially modulate the behavior of Cas10: tag-proximal mismatches depopulate Cas10 from its active state, whereas tag-distal ones yield wildtype-like dynamics. As a result, type III CRISPR systems exhibit broad target specificity and can tolerate target mutations to varying degrees.

### Discrimination Between Self and Nonself

Unlike type I (Cascade), II (Cas9), and V (Cpf1) systems that recognize specific PAM sequences to license foreign DNA, type III systems recognize specific protospacer flanking sequences—ones that are complementary to the crRNA tag—to identify self RNA (Mohanraju et al., 2016). Any noncomplementary flanking sequence will elicit an immune response. The molecular underpinning of this unique discrimination mechanism has remained puzzling, because the type III machinery displays nondiscriminatory binding and cleavage activities on any RNA that contains the protospacer sequence—self or nonself. Here we show that the discriminatory step is manifested in the distinctive Cas10 conformational dynamics. Foreign RNA binding enables Cas10 to quickly sample a large conformational space, including a DNase-active configuration. Engagement with DNA substrates further enhances Cas10’s propensity for residing at the active configuration. On the other hand, self RNA dramatically represses the structural fluctuations of Cas10 and locks it in an inactive configuration. As a result, Cas10 fails to access the DNase-competent state, thereby effectively preventing self-targeting.

Notably, Cas10 harbors robust ssDNA exonucleolytic activity by itself (Ramia et al., 2014), but loses its DNase activity in the apo Cas10-Csm complex (Kazlauskiene et al., 2016). Target RNA binding induces a rotation of Cas10 relative to the rest of the complex in a Cas10-Cmr structure (Taylor et al., 2015). High-resolution structural studies are needed to provide atomic details of the self-RNA-bound Cas10-Csm complex and explain how base pairing between the crRNA tag and the target RNA 3’ flanking sequence deactivates Cas10. Whether degradation of the self RNA transcribed from the host’s CRISPR loci—a consequence of this unique discrimination mechanism—bears any functional impact remains an open question.

### Mutation Tolerance

Phages are constantly evolving to avoid elimination by prokaryotic defense systems. Type I and II CRISPR-Cas systems are extremely sensitive to mutations in the PAM-proximal region of the protospacer, also known as the seed region (Semenova et al., 2011; Wiedenheft et al., 2011). Single-nucleotide mutations in the seed or PAM abolish immunity and cause viral escape. Mismatches in PAM-distal regions are tolerated to some extent but still cause compromised immune responses (Wu et al., 2014). Such strict sequence requirements are related to the process of target recognition, in which initial PAM binding leads to directional unwinding of the dsDNA and formation of the RNA/DNA heteroduplex (R-loop) from the seed throughout the protospacer. Target mutations inhibit R-loop formation and reduce its stability, thereby compromising the efficiency of CRISPR interference (Blosser et al., 2015; Szczelkun et al., 2014). In comparison, the entire single-stranded RNA target sequence in type III systems is directly available for base pairing with crRNA, circumventing the need for duplex unwinding. Hence any single point mutation is unlikely to significantly affect the affinity between the target RNA and the effector complex.

Nonetheless, mismatches do affect type III interference efficiency to varying degrees (Pyenson et al., 2017). Our data offer an explanation for this effect: mismatches in different regions of the protospacer differentially alter the conformational distribution of Cas10. For the *gp43* spacer used in this work, mismatches from tag-distal to tag-proximal regions increasingly populate Cas10 in the inactive mode, resulting in decreasing immunity observed *in vivo*. Based on the few spacers examined in our current and prior studies (Pyenson et al., 2017), this spatial pattern of mismatch sensitivity seems to be generalizable. Nonetheless the quantitative level of inhibition by mismatches is likely spacer specific.

The base-pairing status of the spacer/protospacer region, which resides within the backbone of the effector complex, allosterically modulates the behavior of Cas10. However, none of the mismatched targets abolishes the DNase-competent population of Cas10 to the same extent as the Anti-tag RNA. Consequently, type III systems display broad tolerance to target mutations. Viral escape necessitates a full deletion of the target (Pyenson et al., 2017) or a specific set of mutations that create a perfect match with the crRNA tag, both of which are rare events. Therefore, type III CRISPR-Cas systems evolved an elegant strategy that provides robust immune responses and greatly limits viral escape, while at the same time effectively avoiding autoimmunity.

### Internal Dynamics of the Effector Complex Dictate CRISPR-Cas Immunity

The striking correlation between the conformational distribution of Cas10 and the strength of type III CRISPR immunity enabled us to assign the active and inactive states of Cas10. We identified one subset of states (G_4_) associated with CRISPR interference by the Cas10-Csm complex. It is worth noting that, besides the DNase activity residing in the HD domain, Cas10 also harbors in its Palm domain another catalytic activity that converts ATP into cyclic oligoadenylates (Kazlauskiene et al., 2017; Niewoehner et al., 2017). This signaling molecule activates the Csm6 RNase for nonspecific RNA degradation, which can contribute to the overall type III-A CRISPR-Cas immune response. Future experiments will allow us to distinguish the different conformational states of Cas10 associated with each of its two activities.

The correlation between internal dynamics and enzymatic activity has also been reported for the type II single-subunit effector Cas9 (Dagdas et al., 2017; Sternberg et al., 2015; Yang et al., 2018). The DNA cleavage activity of Cas9 scales with the fraction of time it spends in the DNase-active state. Target mismatches prevent transition from a checkpoint state to the active state. These parallels between Cas9 and Cas10 imply that they are common features for CRISPR-Cas systems.

In summary, we showed that the conformational fluctuations of Cas10 are exquisitely regulated by the complementarity between the target RNA and crRNA. As such, Cas10’s activity is controlled as a gradual dimmer rather than an on-off switch, allowing the host to tune its immune response to an optimal level according to the particular circumstance. The single-molecule imaging platform established here can be used to study Cas10-Csm/Cmr complexes from other species. It will also be interesting to directly observe the concerted action of the transcription complex and the type III CRISPR machinery.

## ACKNOWLEDGMENTS

We thank Poulami Samai, Wenyan Jiang and Rachel Leicher for experimental help, Scott Blanchard (Weill Cornell Medicine) and Taekjip Ha (Johns Hopkins University) for sharing analysis software. L.W. is supported by a Rockefeller University Women & Science postdoctoral fellowship. C.Y.M. is supported by an NIH NRSA postdoctoral fellowship (1F32GM128271-01). M.R.W. is supported by a Rockefeller University Anderson Cancer Center postdoctoral fellowship. J.T.R. is supported by a Boehringer Ingelheim Fonds PhD fellowship. L.A.M. is supported by a Burroughs Wellcome Fund PATH Award, an NIH Director’s Pioneer Award (DP1GM128184) and an HHMI-Simons Faculty Scholar Award. S.L. is supported by the Robertson Foundation, the Quadrivium Foundation, a Monique Weill-Caulier Career Award, a Basil O’Connor Starter Scholar Award from March of Dimes (#5- FY17-61), a Kimmel Scholar Award, and an NIH Pathway to Independence Award (R00GM107365).

## MATERIALS AND METHODS

### Bacterial strains and growth conditions

*Staphylococcus aureus* RN4220 (Kreiswirth et al., 1983) was cultured on Bovine Heart Infusion (BHI) agar plates containing 10 μg/mL erythromycin and 10 μg/mL chloramphenicol to ensure pTarget and pCRISPR plasmid maintenance, respectively. When appropriate, anhydrotetracycline (aTc) was used at a concentration of 0.25 μg/mL to initiate transcription from the P_tet_ promoter.

All expression vectors were transformed into *E. coli* BL21 (DE3) Rosetta 2 cells grown in Terrific Broth medium (Fisher Scientific) containing 100 μg/mL ampicillin and 34 μg/mL chloramphenicol at 37 °C, induced at mid-log phase with 0.5 mM IPTG, and then transferred to 16 °C for overnight expression.

### Plasmid construction

#### Plasmid for heterologous expression of the Cas10-Csm complex in E. coli

pAS1 was constructed based on the plasmid pPS22 harboring the repeat-spacer array and *csm* genes encoding the Csm proteins as well as the processing enzyme Cas6 (Hatoum-Aslan et al., 2013), but modified so that it contains one single spacer targeting the *gp43* gene. To generate the Csm3^D32A^ mutation, pPS86 (Samai et al., 2015) and pAS1 were used as PCR templates with two sets of primers LW10F/R and LW11F/R (see Table S1 for sequences) respectively. The PCR products were joined by Gibson assembly and the mutation was confirmed by DNA sequencing. To attach a SNAP-tag to the N-terminus of Cas10 (or Csm5, Csm4), pSNAP-tag vector (New England Biolabs) and pAS1 were used as PCR templates with primers LW1F/R (or LW7F/R, LW34F/R) and LW2F/R (or LW6F/R, LW35F/R) respectively, and joined by Gibson assembly.

#### Plasmid for transformation assay in Staphylococcus

pCRISPR and pCRISPR∆spc were obtained previously (Hatoum-Aslan et al., 2013; Samai et al., 2015). To clone the pTarget^WT^ plasmid (pJTR48), a DNA fragment containing a gp43 protospacer (Jiang et al., 2016) surrounded by transcriptional terminators on either side (BBa_1006 and BBa_K864501 from http://parts.igem.org/Main_Page) was synthesized by Genewiz (South Plainfield, NJ, USA). This was amplified by PCR using oligos JTR234 and JTR235, and restriction cloned into pE194 (Horinouchi and Weisblum, 1982) amplified with JTR232 and JTR233 and digested with EcoRI and NotI, creating pJTR41. An NheI site was inserted upstream of the protospacer by amplifying pJTR41 with JTR248 and JTR249 by PCR, and ligating the resulting product, and pJTR41, after digestion with NotI and HindIII, creating pJTR43. The aTc-inducible promoter was PCR amplified from pWJ153 (Goldberg et al., 2014) using JTR250 and JTR251, digested with NotI and EcoRI, and ligated upstream of the protospacer with digested pJTR43, creating pJTR46. To add the tetracycline repressor, pWJ153 was PCR amplified using JTR258 and JTR259. The resulting product, and pJTR46, was digested with NheI and HindIII, and the resulting fragments joined by ligation.

To generate mutant pTarget plasmids, mutations in the protospacer sequence or the flanking sequences were introduced via oligonucleotide cassette-based mutagenesis. The pJTR48 plasmid was digested with two restriction enzymes, MfeI and HindIII, flanking the target gp43 sequence. The digested plasmid was then treated with Calf Intestinal Phosphatase (New England Biolabs) for 1 hour at 37 °C, before being purified via a standard spin column DNA cleanup procedure. Oligos (CYM339/340, CYM343/344, CYM372/373, or CYM374/375) containing the mutations (Anti-tag, MM1-10, MM11-20, or MM26-35) (see Table S1) were annealed in a thermocycler, phosphorylated with T4 Polynucleotide Kinase (New England Biolabs) for 1 hour at 37 °C, and spin column purified as well. Digested plasmid and annealed oligo cassettes were then ligated with T4 DNA ligase at 16 °C for 16 hours.

### Protein expression and purification

pAS1 was transformed into *E. coli* BL21 (DE3) Rosetta 2 cells (Merck Millipore), grown in Terrific Broth medium (Fisher Scientific) containing 100 μg/mL ampicillin and 34 μg/mL chloramphenicol at 37 °C until *A*_600_ reached 0.6. Cells were grown for another 16 hours with 0.5 mM IPTG at 16 °C and then harvested by centrifugation. The pellets were resuspended in lysis buffer (50 mM Tris-HCl pH 7.5, 350 mM NaCl, 10 mM imidazole, 1 mM β- mercaptoethanol, 0.1% Triton X-100). The lysate was sonicated and the supernatant was bound to Ni-NTA agarose (Qiagen) in lysis buffer, followed by wash flow using lysis buffer containing 50 mM, 75 mM, and 100 mM imidazole in a stepwise manner. Cas10-Csm complexes loaded with mature crRNA were finally eluted from the Ni-NTA column with lysis buffer containing 250 mM imidazole and subsequently purified on a 1-mL Resource Q column (GE Healthcare) with a linear gradient of 50-1000 mM NaCl. The peak fraction from the Resource Q column was further purified by size exclusion chromatography using Superdex 200 10/300 GL (GE Healthcare) in storage buffer (50 mM Tris-HCl pH 7.5, 150 mM NaCl, 5% glycerol). The mutant (Csm3^D32A^) and SNAP-tagged protein complexes were purified using the same procedure.

### Site-specific fluorescent labeling

#### Protein labeling

SNAP-tagged Cas10-Csm protein complexes were labeled at a concentration of 5 μM with 10 μM SNAP-Surface AlexaFluor647 (New England Biolabs) in 50 mM Tris-HCl pH 7.5, 150 mM NaCl, 5% glycerol, and 1 mM DTT. Labeling mixture was incubated in dark for 2 hours at room temperature. Free dyes were removed by Superdex 200 10/300 GL (GE Healthcare). The labeling efficiency was estimated by measuring *A*_280_ and *A*_650_ with a Nanodrop spectrophotometer (Thermo Fisher Scientific). Labeled samples were flash-frozen in liquid nitrogen and stored in small aliquots at -80 °C.

#### Nucleic acid labeling

DNA and RNA oligonucleotides were purchased from IDT. RNA with a 5’ or 3’ amino modifier was dissolved in 0.1 mL of 0.5 M NaCl, flown through a Sephadex G-25 desalting column (GE Healthcare) to remove traces of ammonia, and then incubated with one pack of Cy3 mono-reactive dye (GE Healthcare) in 0.1 M NaHCO_3_ pH 8.5 at room temperature for 2 hours. Free dyes were removed by Sephadex G-25. Labeled RNAs were subjected to ethanol precipitation and stored in TE buffer (10 mM Tris-HCl pH 8.0, 1 mM EDTA). The labeling efficiency was estimated by measuring *A*_260_ and *A*_550_.

### Bulk RNA cleavage assay

RNA cleavage reactions were performed at room temperature with 20 nM Cy3-labeled RNA and 100 nM Cas10-Csm complexes in a buffer consisting of 50 mM Tris-HCl pH 7.5, 2 mM TCEP, and 0.1 mg/mL BSA. Reaction was initiated by the addition of 10 mM MgCl_2_. Products were collected at time intervals, quenched with 2× loading buffer (90% formamide, 50 mM EDTA, 5% glycerol, 0.1% bromophenol blue), separated on a 12% denaturing polyacrylamide gel, and visualized on a Typhoon imager (GE Healthcare).

### Bacterial transformation assay

*Staphylococcus aureus* RN4220 strains were first transformed with pTarget plasmids carrying the various target sequences. 100 ng of dialyzed plasmid DNA was electroporated into electrocompetent RN4220 cells using a GenePulser Xcell (BioRad) with the following parameters: 2900 V, 25 mF, 100 V, 2 mm. Electroporated strains were then immediately resuspended in 500 μL of Bovine Heart Infusion (BHI) broth and grown at 37 °C with shaking (220 rpm) for 1 hour. Cells were then plated onto BHI agar plates containing 10 μg/mL of erythromycin and left to incubate at 37 °C for 16 hours. Single colonies from the plates were then picked to generate electrocompetent cells carrying the pTarget plasmids. RN4220 strains carrying the pTarget plasmids were then transformed with 100 ng of either pCRISPR or pCRISPRΔspc via the same electroporation protocol as described above. Following growth at 37 °C in BHI broth for 1 hour, cells were spun down on a table-top centrifuge at 6000 rpm for 3 minutes and resuspended in 1 mL of fresh BHI broth. 100 μL of the resuspended culture was plated onto BHI agar plates with 10 μg/mL of erythromycin and 10 μg/mL of chloramphenicol; another 100 μL of the same culture was plated onto BHI agar plates with 10 μg/mL of erythromycin, 10 μg/mL of chloramphenicol, and 0.25 μg/mL of anhydrotetracycline (aTc). Plates without aTc were incubated at 37 °C for 24 hours, while plates with aTc were incubated for 48 hours. For each plate, the colony forming units per μg of plasmid (CFU/μg) was calculated. To quantify the efficiency of targeting for each transformed culture, the CFUs in the presence of aTc was divided by the CFUs in the absence of aTc.

### Single-molecule experiments

#### Data acquisition

Single-molecule experiments were performed at room temperature (23 ± 1 °C) in an imaging buffer consisting of 50 mM Tris-HCl pH 7.5, 2 mM TCEP, 0.1 mg/mL BSA, 1 mM EDTA or 10 mM MgCl_2_, and an oxygen scavenging system containing 1% w/v D-glucose, 1 mg/mL glucose oxidase (Sigma-Aldrich), 0.04 mg/mL catalase (Sigma-Aldrich) and 2 mM Trolox (Sigma-Aldrich). The microfluidic flow chambers were passivated with a mixture of polyethylene glycol (PEG) and biotin-PEG (Laysan Bio), incubated with 40 μL of 0.1 mg/mL streptavidin (Thermo Fisher Scientific), and washed with 100 μL of T50 (10 mM Tris-HCl pH 8.0, 50 mM NaCl). Then 40 μL of 500 pM biotinylated RNA molecules were injected into the chamber and immobilized through biotin-streptavidin linkage. 40 μL of 10 nM labeled Cas10-Csm complexes were added to the chamber and incubated for 5 minutes before imaging. Donor and acceptor fluorescence signals were collected on a total-internal-reflection fluorescence microscope (Olympus IX83 cellTIRF) and detected by an EMCCD camera (Andor iXon Ultra897) with a frame rate of 300 ms.

#### Data analysis

Fluorescence time trajectories of individual RNA molecules were extracted and analyzed by the SPARTAN software (Juette et al., 2016). The FRET value was calculated as *I*_A_/(*I*_D_+*I*_A_), where *I*_D_ and *I*_A_ represent the donor and acceptor fluorescence intensity, respectively. Dynamic FRET traces were analyzed by a hidden-Markov-model based software HaMMy (McKinney et al., 2006). FRET contour plots and histograms were built from at least 600 molecules from multiple fields of imaging and plotted by Origin (OriginLab). Transition density plots were generated using a custom code written in MATLAB.

### Quantification and statistical analysis

Statistical significance was determined by unpaired two-tailed Student’s t tests using GraphPad Prism 7. The difference between two groups was considered statistically significant when the p value is less than 0.05 (^*^, p < 0.05; ^**^, p < 0.01; ^***^, p < 0.001; ns, not significant). The number of molecules analyzed or experiments repeated is indicated in the figure legends.

---

**Figure S1.**
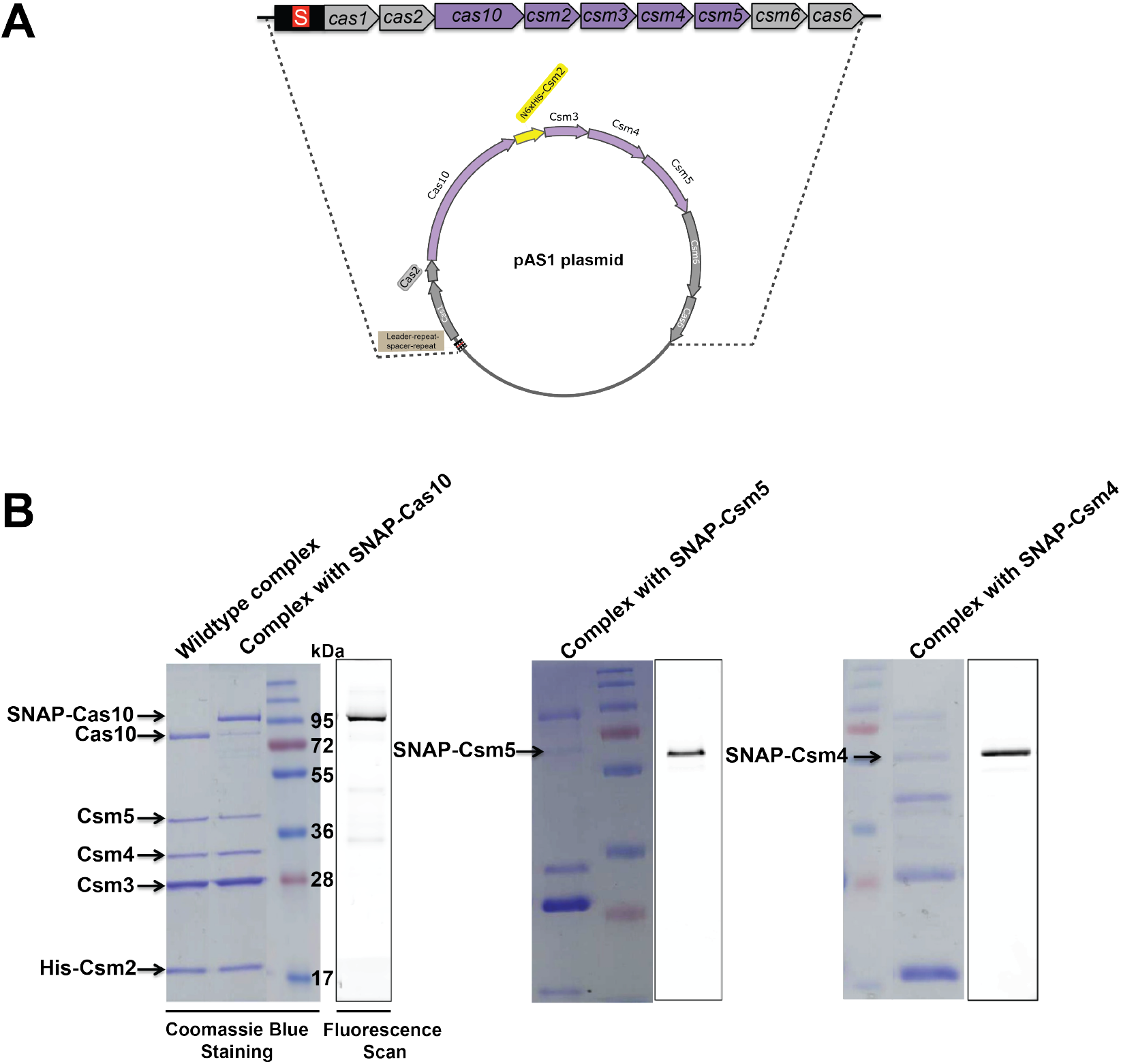
Cas10-Csm Complex Purification and Site-specific Labeling. (**A**) Schematic of the plasmid that encodes an *S. epidermidis* type III-A CRISPR-Cas locus containing one single spacer targeting the *gp43* gene of phage ΦNM1γ6. (**B**) SDS-PAGE analysis of purified wildtype and SNAP-tagged Cas10-Csm complexes. Fluorescence scans show specific labeling of the SNAP-tagged subunit (Cas10, Csm5 and Csm4 in the left, middle and right gel, respectively).

**Figure S2.**
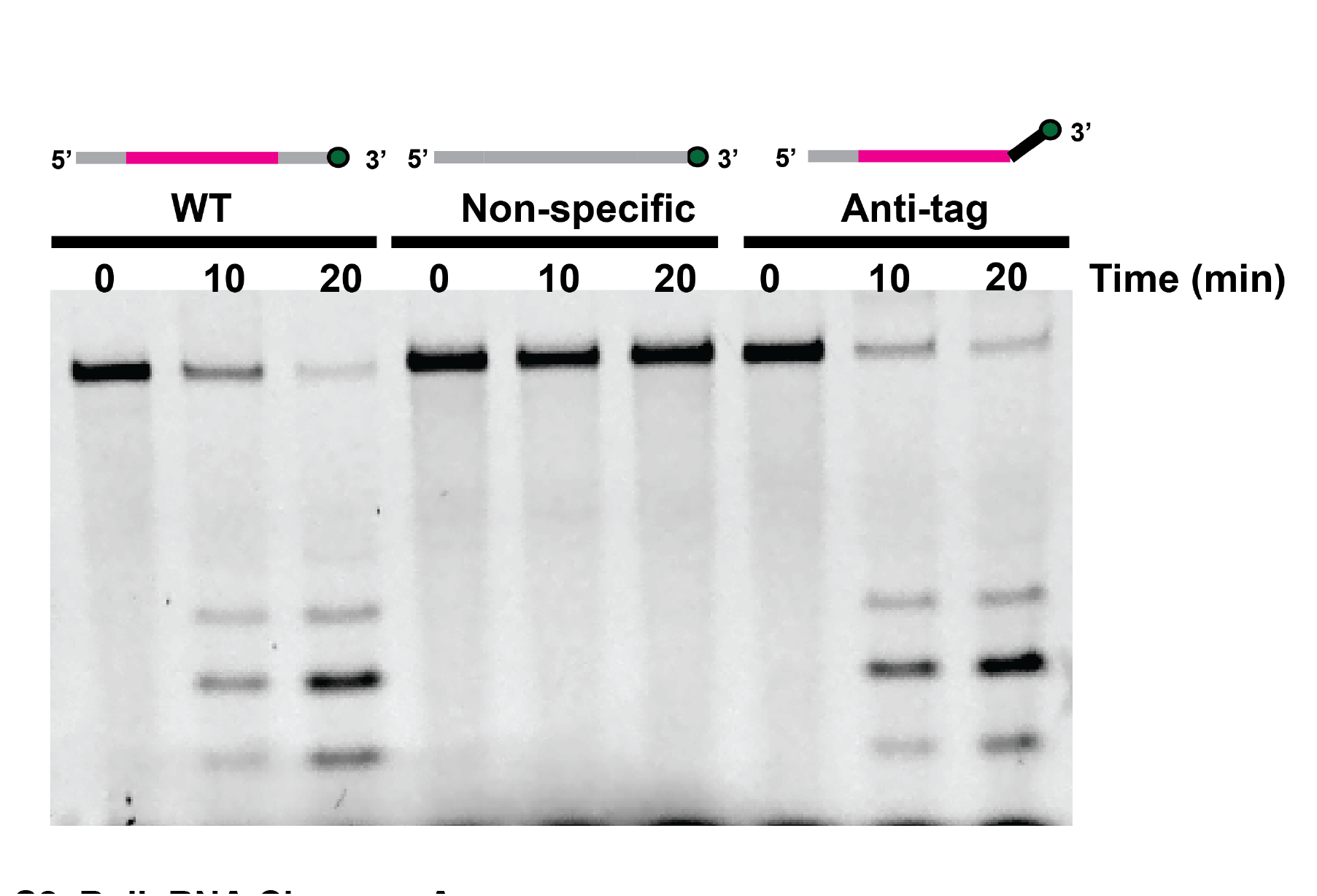
Bulk RNA Cleavage Assay. Shown is a comparison of RNA cleavage activities of the Cas10-Csm complex on WT, Anti-tag and Non-specific RNAs. The 6-nucleotide periodicity due to cleavage by the multiple copies of Csm3 was observed.

**Figure S3.**
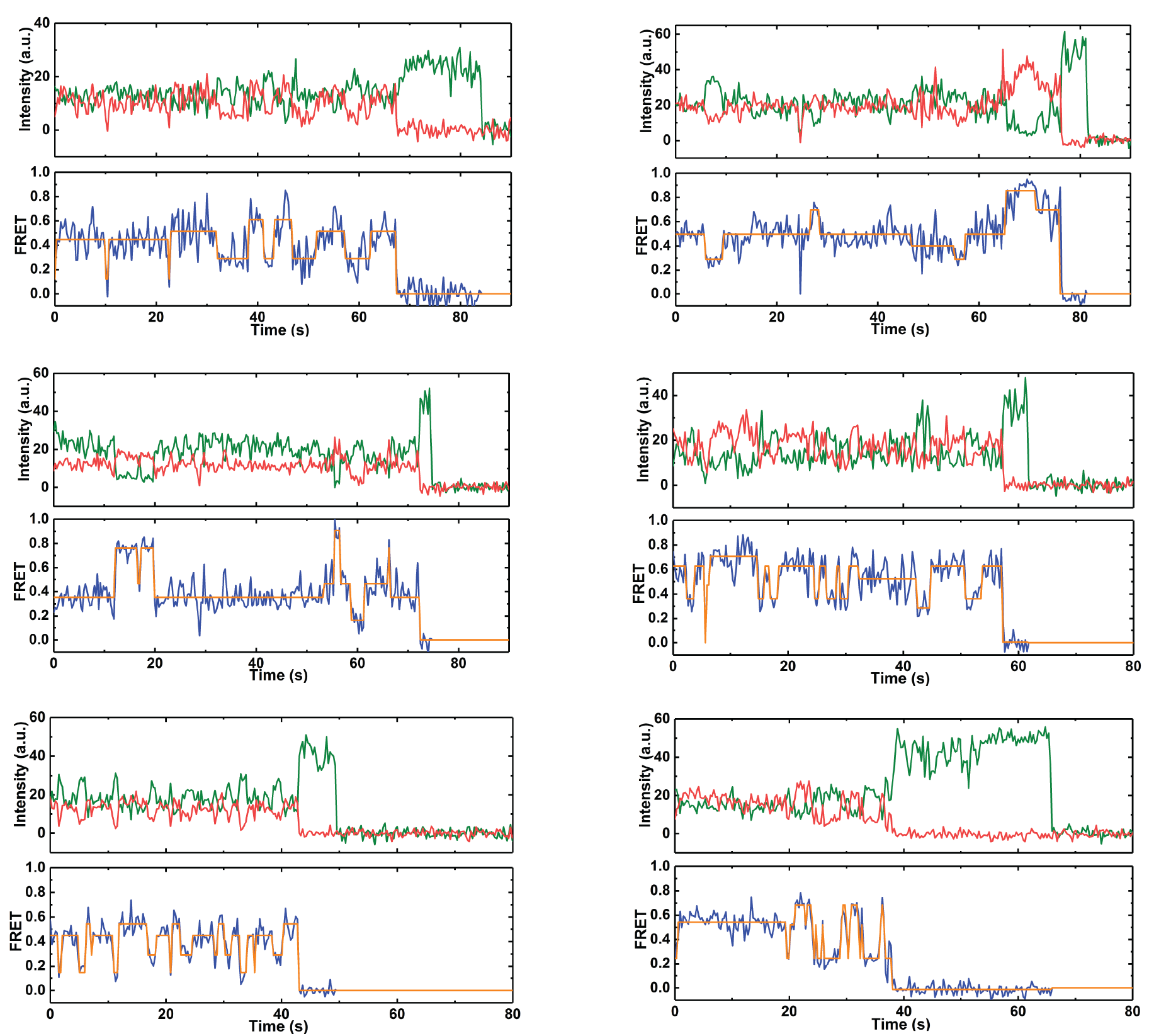
Additional Representative Fluorescence and FRET Traces Using AlexaFluor647-Labeled Cas10 and Cy3-Labeled WT RNA. Idealized FRET states from hidden-Markov-modeling (HMM) analysis are overlaid in orange.

**Figure S4.**
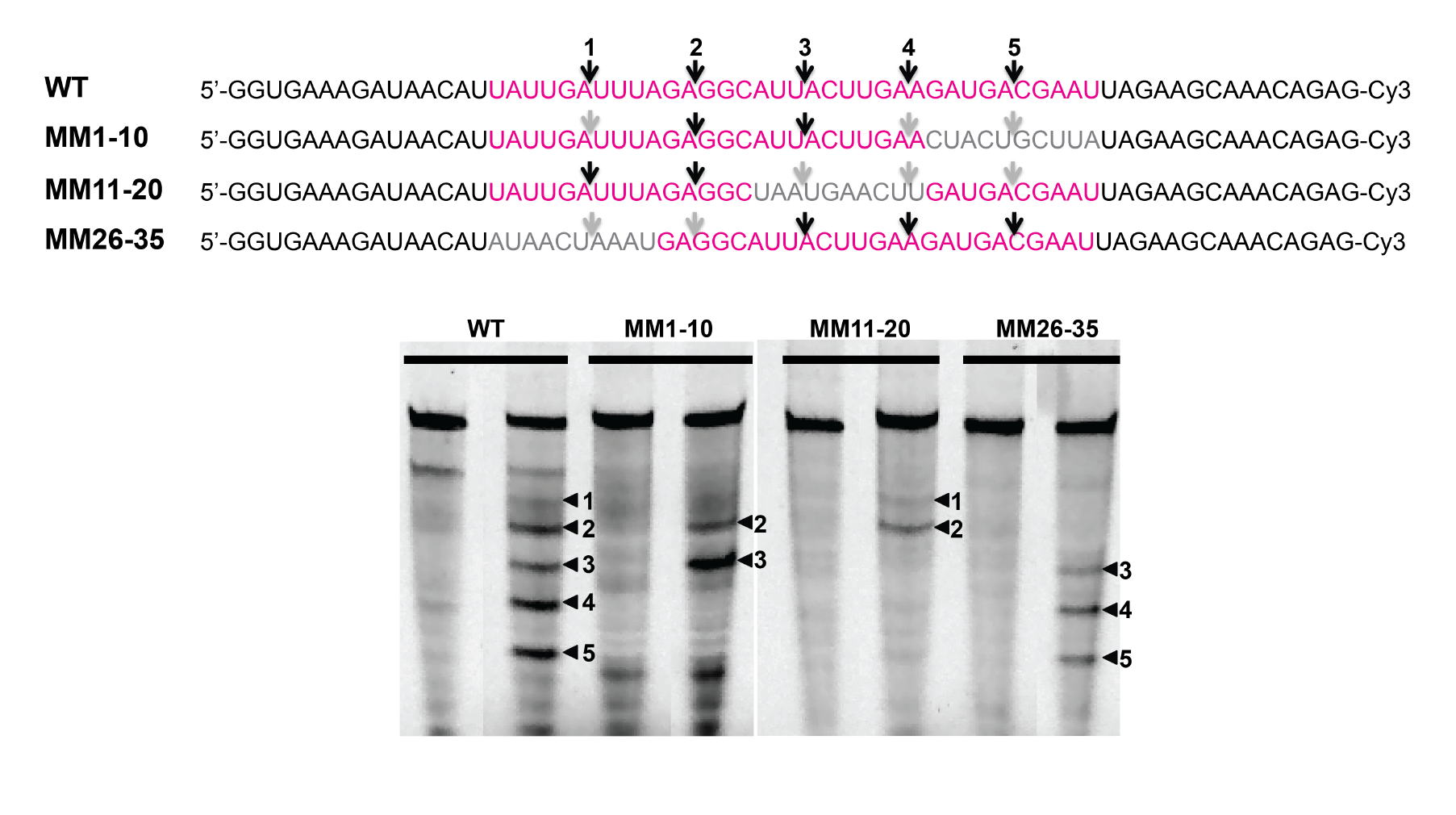
Bulk RNA Cleavage Assay with Mismatched RNA Targets. Black arrows indicate cleavage sites. Grey arrows indicate abolished cleavage sites due to target mutation.

**Figure S5.**
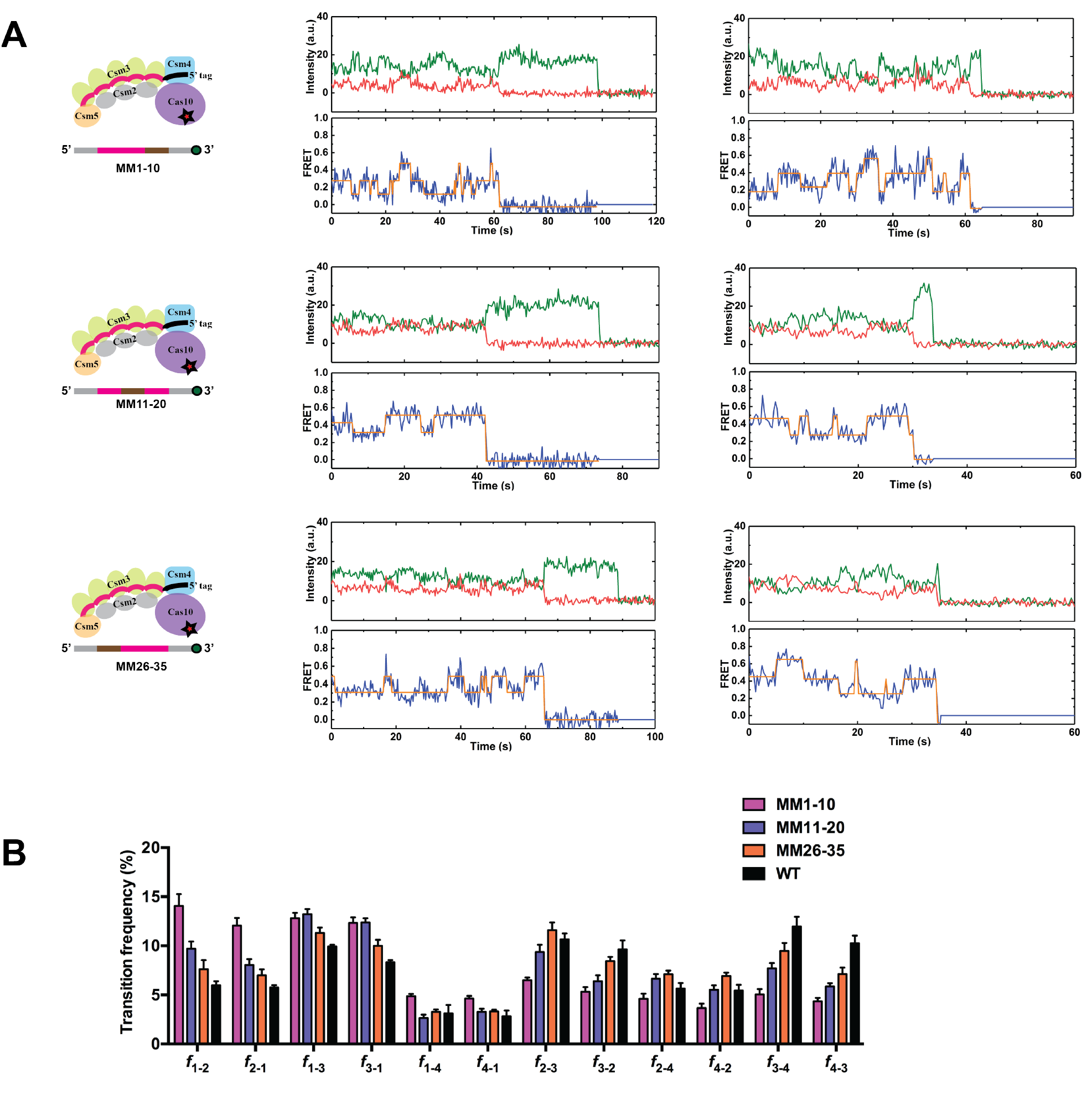
Analyses of the Dynamic Interactions between the Cas10-Csm Complex and Mismatched RNA Targets. (**A**) Representative fluorescence and FRET traces for mismatched RNA targets (MM1-10, MM11-20 and MM26-35). FRET donor and acceptor were placed on the 3’ end of RNA and Cas10, respectively. Idealized FRET states from HMM analysis are overlaid in orange. (**B**) Transition frequencies between different FRET groups for wildtype and mutant RNA targets. For example, *f*_1-2_ denotes the transition frequency from G_1_ states to G_2_ states. Data are represented as mean ± SEM.

**Figure S6.**
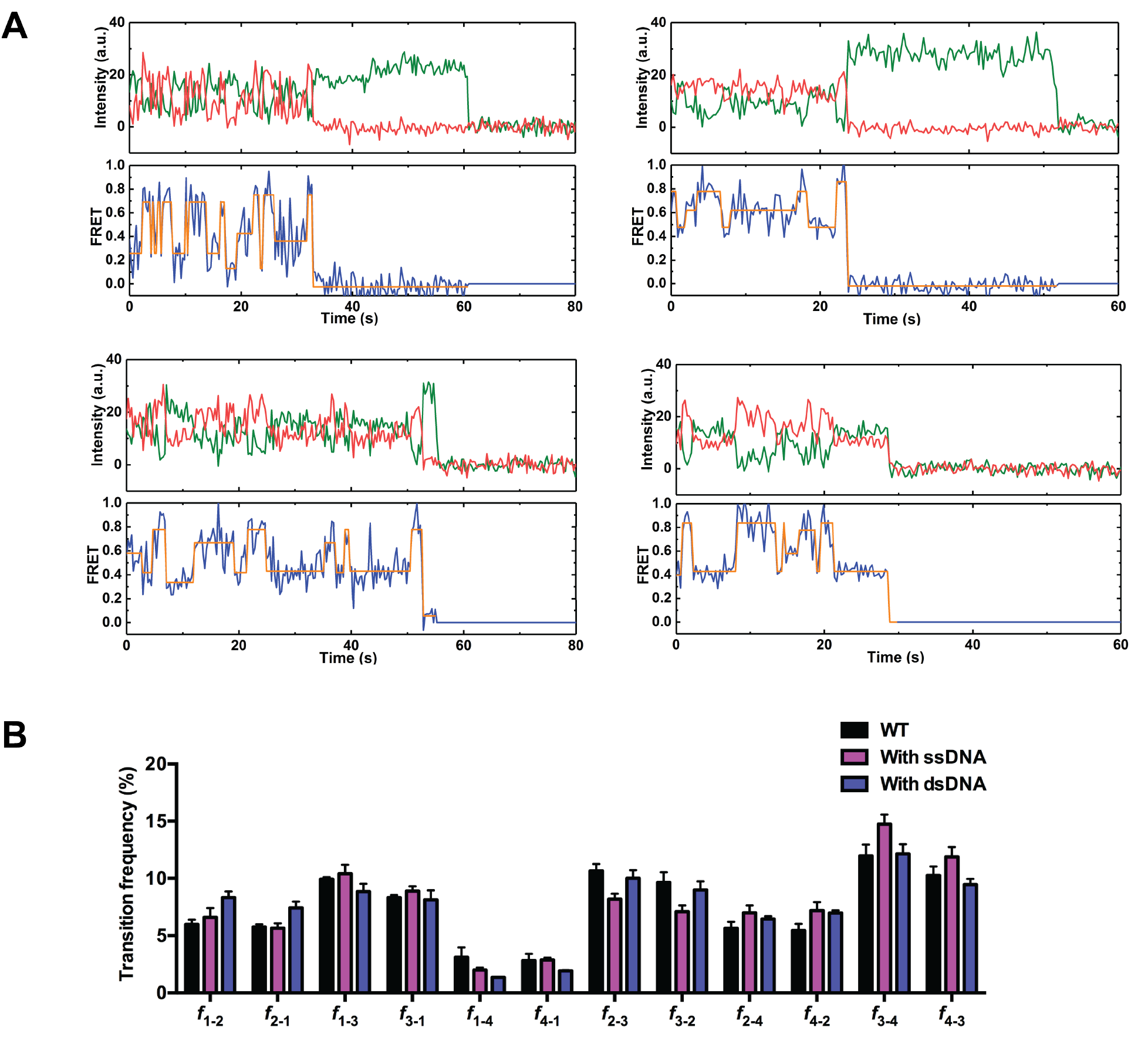
Effects of DNA on the Behavior of Cas10. (**A**) Additional representative fluorescence and FRET traces of the Cas10-Csm complex interacting with the WT RNA target in the presence of 500 nM ssDNA in solution. Idealized FRET states from HMM analysis are overlaid in orange. FRET donor and acceptor were placed on the 3’ end of RNA and Cas10, respectively. (**B**) Transition frequencies between different FRET groups for the WT RNA in the absence and presence of ssDNA or dsDNA. Data are represented as mean ± SEM.

**Table S1.**
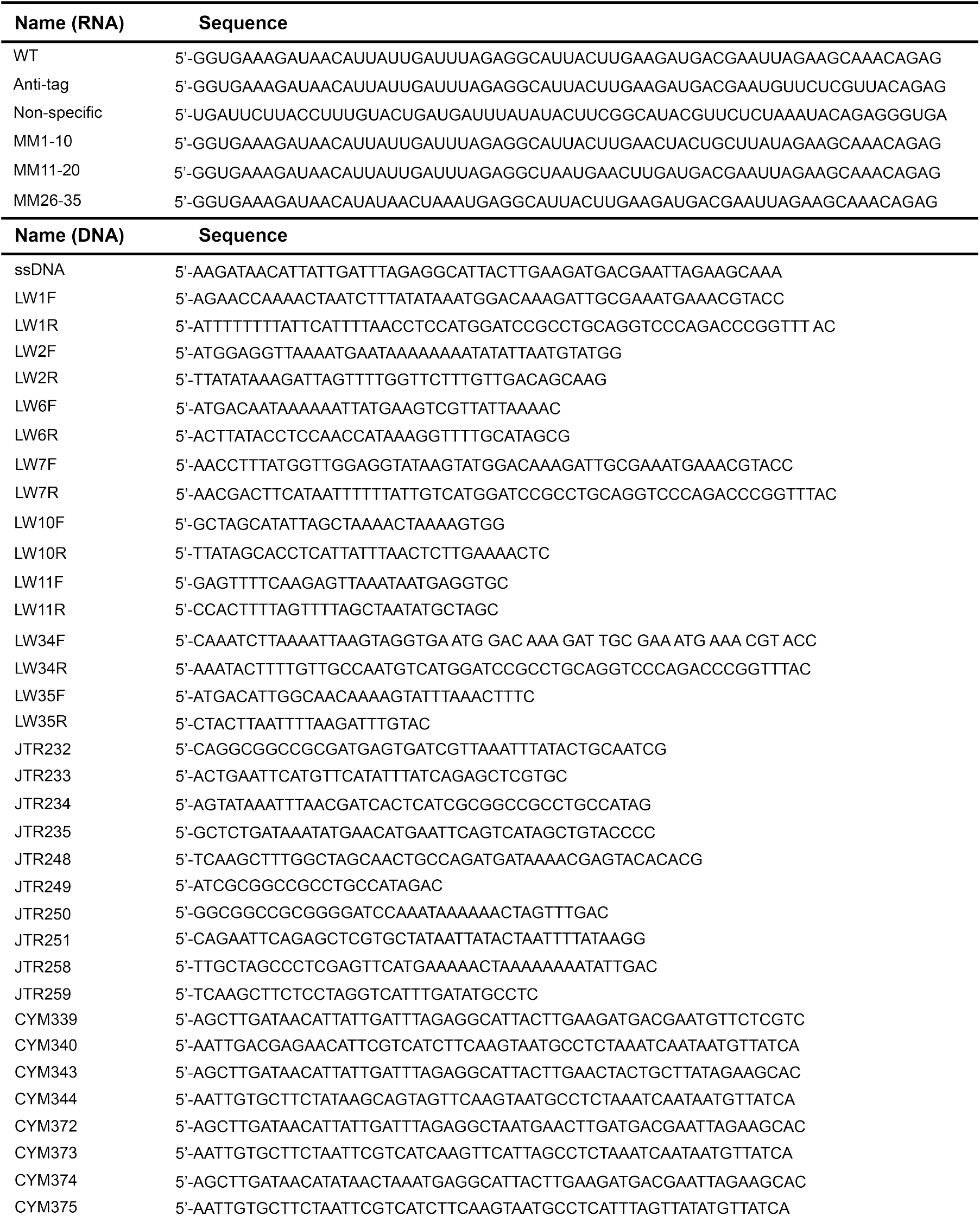
Sequences of Oligonucleotides Used in This Study.

## REFERENCES

Barrangou, R., and Marraffini, L.A. (2014. CRISPR-Cas systems: Prokaryotes upgrade to adaptive immunity. Molecular cell 54, 234–244.

Blosser, T.R., Loeff, L., Westra, E.R., Vlot, M., Kunne, T., Sobota, M., Dekker, C., Brouns, S.J.J., and Joo, C. (2015. Two distinct DNA binding modes guide dual roles of a CRISPR-Cas protein complex. Molecular cell 58, 60–70.

Boehm, T. (2006. Quality control in self/nonself discrimination. Cell 125, 845–858.

Dagdas, Y.S., Chen, J.S., Sternberg, S.H., Doudna, J.A., and Yildiz, A. (2017. A conformational checkpoint between DNA binding and cleavage by CRISPR-Cas9. Science advances 3, eaao0027.

Deng, L., Garrett, R.A., Shah, S.A., Peng, X., and She, Q. (2013. A novel interference mechanism by a type IIIB CRISPR-Cmr module in Sulfolobus. Molecular microbiology 87, 1088–1099.

Elmore, J.R., Sheppard, N.F., Ramia, N., Deighan, T., Li, H., Terns, R.M., and Terns, M.P. (2016. Bipartite recognition of target RNAs activates DNA cleavage by the Type III-B CRISPR-Cas system. Genes & development 30, 447–459.

Estrella, M.A., Kuo, F.T., and Bailey, S. (2016. RNA-activated DNA cleavage by the Type III-B CRISPR-Cas effector complex. Genes & development 30, 460–470.

Gasiunas, G., Barrangou, R., Horvath, P., and Siksnys, V. (2012. Cas9-crRNA ribonucleoprotein complex mediates specific DNA cleavage for adaptive immunity in bacteria. Proceedings of the National Academy of Sciences of the United States of America 109, E2579–2586.

Goldberg, G.W., Jiang, W., Bikard, D., and Marraffini, L.A. (2014. Conditional tolerance of temperate phages via transcription-dependent CRISPR-Cas targeting. Nature 514, 633–637.

Hale, C.R., Zhao, P., Olson, S., Duff, M.O., Graveley, B.R., Wells, L., Terns, R.M., and Terns, M.P. (2009. RNA-guided RNA cleavage by a CRISPR RNA-Cas protein complex. Cell 139, 945–956.

Hatoum-Aslan, A., Samai, P., Maniv, I., Jiang, W., and Marraffini, L.A. (2013. A ruler protein in a complex for antiviral defense determines the length of small interfering CRISPR RNAs. The Journal of biological chemistry 288, 27888–27897.

Horinouchi, S., and Weisblum, B. (1982. Nucleotide sequence and functional map of pE194, a plasmid that specifies inducible resistance to macrolide, lincosamide, and streptogramin type B antibodies. Journal of bacteriology 150, 804–814.

Jiang, W., Samai, P., and Marraffini, L.A. (2016. Degradation of Phage Transcripts by CRISPR-Associated RNases Enables Type III CRISPR-Cas Immunity. Cell 164, 710–721.

Josephs, E.A., Kocak, D.D., Fitzgibbon, C.J., McMenemy, J., Gersbach, C.A., and Marszalek, P.E. (2015. Structure and specificity of the RNA-guided endonuclease Cas9 during DNA interrogation, target binding and cleavage. Nucleic acids research 43, 8924–8941.

Juette, M.F., Terry, D.S., Wasserman, M.R., Altman, R.B., Zhou, Z., Zhao, H., and Blanchard, S.C. (2016. Single-molecule imaging of non-equilibrium molecular ensembles on the millisecond timescale. Nature methods 13, 341–344.

Jung, C., Hawkins, J.A., Jones, S.K.Jr.,, Xiao, Y., Rybarski, J.R., Dillard, K.E., Hussmann, J., Saifuddin, F.A., Savran, C.A., Ellington, A.D., et al. (2017. Massively Parallel Biophysical Analysis of CRISPR-Cas Complexes on Next Generation Sequencing Chips. Cell 170, 35–47 e13.

Kazlauskiene, M., Kostiuk, G., Venclovas, C., Tamulaitis, G., and Siksnys, V. (2017. A cyclic oligonucleotide signaling pathway in type III CRISPR-Cas systems. Science 357, 605–609.

Kazlauskiene, M., Tamulaitis, G., Kostiuk, G., Venclovas, C., and Siksnys, V. (2016. Spatiotemporal Control of Type III-A CRISPR-Cas Immunity: Coupling DNA Degradation with the Target RNA Recognition. Molecular cell 62, 295–306.

Koonin, E.V., Makarova, K.S., and Zhang, F. (2017. Diversity, classification and evolution of CRISPR-Cas systems. Current opinion in microbiology 37, 67–78.

Kreiswirth, B.N., Lofdahl, S., Betley, M.J., O’Reilly, M., Schlievert, P.M., Bergdoll, M.S., and Novick, R.P. (1983. The toxic shock syndrome exotoxin structural gene is not detectably transmitted by a prophage. Nature 305, 709–712.

Lim, Y., Bak, S.Y., Sung, K., Jeong, E., Lee, S.H., Kim, J.S., Bae, S., and Kim, S.K. (2016. Structural roles of guide RNAs in the nuclease activity of Cas9 endonuclease. Nature communications 7, 13350.

Loeff, L., Brouns, S.J.J., and Joo, C. (2018. Repetitive DNA Reeling by the Cascade-Cas3 Complex in Nucleotide Unwinding Steps. Molecular cell 70, 385–394 e383.

Manica, A., Zebec, Z., Steinkellner, J., and Schleper, C. (2013. Unexpectedly broad target recognition of the CRISPR-mediated virus defence system in the archaeon Sulfolobus solfataricus. Nucleic acids research 41, 10509–10517.

Maniv, I., Jiang, W., Bikard, D., and Marraffini, L.A. (2016. Impact of Different Target Sequences on Type III CRISPR-Cas Immunity. Journal of bacteriology 198, 941–950.

Marraffini, L.A., and Sontheimer, E.J. (2010. Self versus non-self discrimination during CRISPR RNA-directed immunity. Nature 463, 568–571.

McKinney, S.A., Joo, C., and Ha, T. (2006. Analysis of single-molecule FRET trajectories using hidden Markov modeling. Biophysical journal 91, 1941–1951.

Mohanraju, P., Makarova, K.S., Zetsche, B., Zhang, F., Koonin, E.V., and van der Oost, J. (2016. Diverse evolutionary roots and mechanistic variations of the CRISPR-Cas systems. Science 353, aad5147.

Mojica, F.J., Diez-Villasenor, C., Garcia-Martinez, J., and Almendros, C. (2009. Short motif sequences determine the targets of the prokaryotic CRISPR defence system. Microbiology 155, 733–740.

Niewoehner, O., Garcia-Doval, C., Rostol, J.T., Berk, C., Schwede, F., Bigler, L., Hall, J., Marraffini, L.A., and Jinek, M. (2017. Type III CRISPR-Cas systems produce cyclic oligoadenylate second messengers. Nature 548, 543–548.

Osawa, T., Inanaga, H., Sato, C., and Numata, T. (2015. Crystal structure of the CRISPR-Cas RNA silencing Cmr complex bound to a target analog. Molecular cell 58, 418–430.

Peng, W., Feng, M., Feng, X., Liang, Y.X., and She, Q. (2015. An archaeal CRISPR type III-B system exhibiting distinctive RNA targeting features and mediating dual RNA and DNA interference. Nucleic acids research 43, 406–417.

Pyenson, N.C., Gayvert, K., Varble, A., Elemento, O., and Marraffini, L.A. (2017. Broad Targeting Specificity during Bacterial Type III CRISPR-Cas Immunity Constrains Viral Escape. Cell host & microbe 22, 343–353 e343.

Pyenson, N.C., and Marraffini, L.A. (2017. Type III CRISPR-Cas systems: when DNA cleavage just isn’t enough. Current opinion in microbiology 37, 150–154.

Ramia, N.F., Tang, L., Cocozaki, A.I., and Li, H. (2014. Staphylococcus epidermidis Csm1 is a 3’-5’ exonuclease. Nucleic acids research 42, 1129–1138.

Redding, S., Sternberg, S.H., Marshall, M., Gibb, B., Bhat, P., Guegler, C.K., Wiedenheft, B., Doudna, J.A., and Greene, E.C. (2015. Surveillance and Processing of Foreign DNA by the Escherichia coli CRISPR-Cas System. Cell 163, 854–865.

Samai, P., Pyenson, N., Jiang, W., Goldberg, G.W., Hatoum-Aslan, A., and Marraffini, L.A. (2015. Co-transcriptional DNA and RNA Cleavage during Type III CRISPR-Cas Immunity. Cell 161, 1164–1174.

Semenova, E., Jore, M.M., Datsenko, K.A., Semenova, A., Westra, E.R., Wanner, B., van der Oost, J., Brouns, S.J., and Severinov, K. (2011. Interference by clustered regularly interspaced short palindromic repeat (CRISPR) RNA is governed by a seed sequence. Proceedings of the National Academy of Sciences of the United States of America 108, 10098–10103.

Singh, D., Sternberg, S.H., Fei, J., Doudna, J.A., and Ha, T. (2016. Real-time observation of DNA recognition and rejection by the RNA-guided endonuclease Cas9. Nature communications 7, 12778.

Staals, R.H., Zhu, Y., Taylor, D.W., Kornfeld, J.E., Sharma, K., Barendregt, A., Koehorst, J.J., Vlot, M., Neupane, N., Varossieau, K., et al. (2014. RNA targeting by the type III-A CRISPR-Cas Csm complex of Thermus thermophilus. Molecular cell 56, 518–530.

Staals, R.H.J., Agari, Y., Maki-Yonekura, S., Zhu, Y., Taylor, D.W., van Duijn, E., Barendregt, A., Vlot, M., Koehorst, J.J., Sakamoto, K., et al. (2013. Structure and activity of the RNA-targeting Type III-B CRISPR-Cas complex of Thermus thermophilus. Molecular cell 52, 135–145.

Sternberg, S.H., LaFrance, B., Kaplan, M., and Doudna, J.A. (2015. Conformational control of DNA target cleavage by CRISPR-Cas9. Nature 527, 110–113.

Sternberg, S.H., Redding, S., Jinek, M., Greene, E.C., and Doudna, J.A. (2014. DNA interrogation by the CRISPR RNA-guided endonuclease Cas9. Nature 507, 62–67.

Szczelkun, M.D., Tikhomirova, M.S., Sinkunas, T., Gasiunas, G., Karvelis, T., Pschera, P., Siksnys, V., and Seidel, R. (2014. Direct observation of R-loop formation by single RNA-guided Cas9 and Cascade effector complexes. Proceedings of the National Academy of Sciences of the United States of America 111, 9798–9803.

Tamulaitis, G., Kazlauskiene, M., Manakova, E., Venclovas, C., Nwokeoji, A.O., Dickman, M.J., Horvath, P., and Siksnys, V. (2014. Programmable RNA shredding by the type III-A CRISPR-Cas system of Streptococcus thermophilus. Molecular cell 56, 506–517.

Tamulaitis, G., Venclovas, C., and Siksnys, V. (2017. Type III CRISPR-Cas Immunity: Major Differences Brushed Aside. Trends in microbiology 25, 49–61.

Taylor, D.W., Zhu, Y., Staals, R.H., Kornfeld, J.E., Shinkai, A., van der Oost, J., Nogales, E., and Doudna, J.A. (2015. Structural biology. Structures of the CRISPR-Cmr complex reveal mode of RNA target positioning. Science 348, 581–585.

van der Oost, J., Westra, E.R., Jackson, R.N., and Wiedenheft, B. (2014. Unravelling the structural and mechanistic basis of CRISPR-Cas systems. Nature reviews Microbiology 12, 479–492.

Wiedenheft, B., van Duijn, E., Bultema, J.B., Waghmare, S.P., Zhou, K., Barendregt, A., Westphal, W., Heck, A.J., Boekema, E.J., Dickman, M.J., et al. (2011. RNA-guided complex from a bacterial immune system enhances target recognition through seed sequence interactions. Proceedings of the National Academy of Sciences of the United States of America 108, 10092–10097.

Wu, X., Kriz, A.J., and Sharp, P.A. (2014. Target specificity of the CRISPR-Cas9 system. Quantitative biology 2, 59–70.

Yang, M., Peng, S., Sun, R., Lin, J., Wang, N., and Chen, C. (2018. The Conformational Dynamics of Cas9 Governing DNA Cleavage Are Revealed by Single-Molecule FRET. Cell reports 22, 372–382.

Zhang, J., Rouillon, C., Kerou, M., Reeks, J., Brugger, K., Graham, S., Reimann, J., Cannone, G., Liu, H., Albers, S.V., et al. (2012. Structure and mechanism of the CMR complex for CRISPR-mediated antiviral immunity. Molecular cell 45, 303–313.

